# Titin switches from an extensible spring to a mechanical rectifier upon muscle activation

**DOI:** 10.1101/2021.08.06.455239

**Authors:** Caterina Squarci, Pasquale Bianco, Massimo Reconditi, Marco Caremani, Theyencheri Narayanan, Andras Málnási Csizmadia, Marco Linari, Vincenzo Lombardi, Gabriella Piazzesi

## Abstract

In contracting striated muscle titin acts as a spring in parallel with the array of myosin motors in each half-sarcomere and could prevent the intrinsic instability of thousands of serially linked half-sarcomeres, if its stiffness, at physiological sarcomere lengths (SL), were ten times larger than reported. Here we define titin mechanical properties during tetanic stimulation of single fibres of frog muscle by suppressing myosin motor responses with Para-Nitro-Blebbistatin, which is able to freeze thick filament in the resting state. We discover that thin filament activation switches I-band titin spring from the large SL-dependent extensibility of the OFF-state to an ON-state in which titin acts as a SL-independent mechanical rectifier, allowing free shortening while opposing stretch with an effective stiffness 4 pN nm^−1^ per half-thick filament. In this way during contraction titin limits weak half-sarcomere elongation to a few % and, also, provides an efficient link for mechanosensing-based thick filament activation.

## INTRODUCTION

Force and shortening in striated muscle are generated by the cyclical ATP-driven interactions of the motor protein myosin II, arranged in two bipolar arrays on the thick filaments originating at the midpoint of each sarcomere (M-line), with the nearby thin, actin-containing filaments originating at the sarcomere extremities (Z-line, Figure 1A). In each half-sarcomere myosin motors are mechanically coupled as parallel force generators and the collective force depends on the number of motors available for actin-attachment and thus on the degree of overlap between thick and thin filaments (Figure 1B, black circles) (Gordon et al., 1966). The half-sarcomere is the basic functional unit in which the emergent properties from the arrays of myosin motors, the interdigitating thin filaments and other regulatory and cytoskeleton proteins (Figure 1C) account for the mechanical performance of muscle and its regulation. Macroscopic production of power during contraction of striated muscle depends on maintenance of the ordered configuration of the half-sarcomere on different hierarchical levels both over the cell cross-section (for force) and through the serial arrangement of thousands of half-sarcomeres (for shortening velocity). However, the presence of variability in the force generated by in series half-sarcomeres results in instability, with lengthening of the weaker half-sarcomeres caused by shortening of the stronger ones (Julian and Morgan, 1979). In the descending limb of the active force–SL relation (Figure 1B, black circles), this phenomenon would progressively increase sarcomere length inhomogeneity, unless in the weaker half-sarcomeres the rising strain in the spring in parallel with myosin motors, constituted by the I-band region of titin, contributed to equilibrate the force.

**Figure 1.**
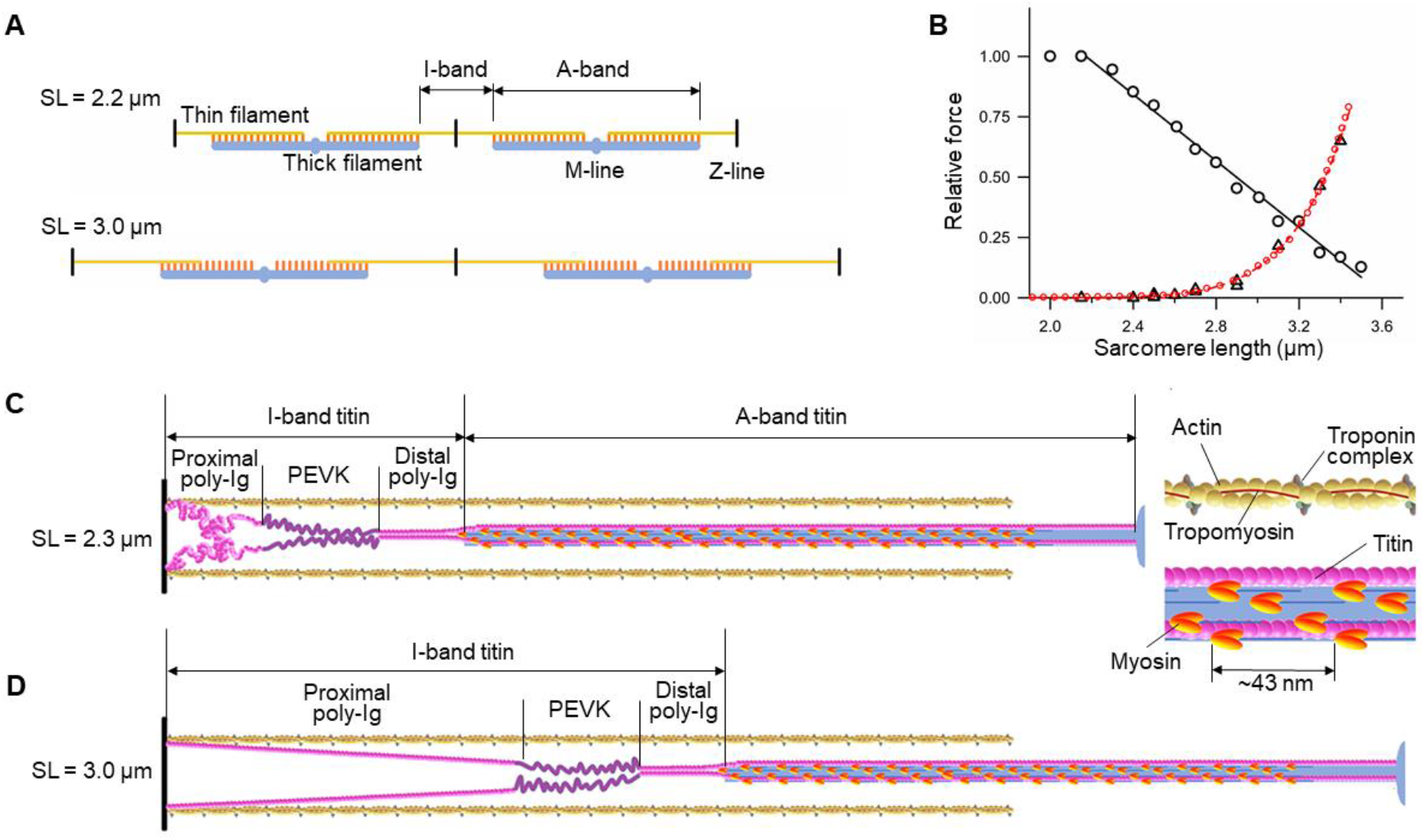
The structure-function of myofilaments and titin in relation to the length of the sarcomere. **A**. Overview of the thick filament (blue), myosin motors (orange), and thin filament (yellow) at SL 2.2 μm (full overlap) and 3.0 μm (partial overlap). **B.** Relation between SL and either active force at the plateau of the isometric tetanic contraction (black circles, fitted by the equation *T*= 2.5 – 0.69×SL in the SL range >2.2 μm, continuous line), or passive force (triangles, fitted by an exponential equation), red dashed line, and by a model with two serially-linked worm like chains, red circles, see also Note S1 and Figure S1). **C.** Protein disposition on the thin and thick filaments in the half-sarcomere at rest at 2.3 μm SL. M-line on the right and Z-line on the left. Inset: overlap region on an enlarged scale. Myosin motors (orange) lie on their tails (blue), tilted toward the M-line; actin monomers form the thin filament (yellow) with tropomyosin (brown) and troponin complex (grey); titin (magenta) in the I-band is constituted by the proximal poly-Ig domain, the PEVK region (dark magenta) and the stiff distal poly-Ig domain. Only two of the six titin molecules per half-thick filament are represented for simplicity. **D.** Straightening of the proximal poly-Ig domain by passive stretch to 3.0 μm SL.

The giant protein titin spans the whole half-sarcomere (Figure 1C, magenta), first through the I-band, connecting the Z-line at the end of the sarcomere with the tip of the myosin filament, and then through the A-band, associated to thick filament (six molecules per thick filament (Trinick, 1996; Whiting et al., 1989)) up to the M-line at the centre of the sarcomere (Labeit and Kolmerer, 1995; Maruyama et al., 1977; Wang et al., 1979). While the A-band region of titin is made inextensible by its tight association to the thick filament, the I-band region is able to transmit the stress also when no myosin motors are attached to actin and thus is responsible for the passive force developed when a skeletal muscle fibre is stretched at rest (Figure 1B, triangles) (Anderson and Granzier, 2012; Furst et al., 1988; Itoh et al., 1988; Linke, 2018; Trombitas et al., 1991). Of the three serially linked spring elements in the I-band titin (Figure 1C), only two account for titin extensibility: the proximal tandem immunoglobulin-like (poly-Ig) domain, and the unique sequence rich in proline, glutamate, valine, and lysine residues (PEVK domain), as the distal poly-Ig-domain forms a much stiffer end-filament composed of the six titin molecules attaching to the tip of the thick filament (Bennett et al., 1997). Both the proximal poly-Ig domain (hereinafter called poly-Ig domain) and the PEVK domain, which both exhibit variable muscle-type specific lengths (Freiburg et al., 2000; Labeit and Kolmerer, 1995), are responsible for the differences in passive force–SL relations(Neagoe et al., 2003).

*In situ* studies using immunofluorescence and immunoelectron microscopy on skinned fibres and myofibrils of mammalian skeletal muscle demonstrated that, at SL < 2.7 μm, the poly-Ig domain is responsible for the large extensibility of the muscle sarcomere by straightening out of randomly bent elements, while at larger SL, where the Ig-domain spring has attained its contour length and passive force increases more steeply, sarcomere extensibility is accounted for by the PEVK domain (Linke et al., 1998a; Linke et al., 1996; Linke et al., 1998b; Trombitas et al., 1998).

In the absence of any contribution from parallel elastic elements of the extra-cellular matrix (ECM), as it is the case of skinned fibres and myofibrils (Linke *et al*., 1998a; Linke *et al*., 1998b; Trombitas *et al*., 1998), but also of intact fibres of frog muscle (Magid and Law, 1985; Meyer and Lieber, 2018), passive force starts to rise at SL > 2.5 μm and attains a significant fraction (≥ 10%) of the active tetanic force at SL ≥ 3 μm (Figure 1B, see also Figure S1) (Granzier and Wang, 1993; Linke *et al*., 1998a; Linke *et al*., 1998b; Reconditi et al., 2014). This indicates that, in the range of physiological SL (classically < 2.7 μm), the underlying static stiffness of titin is too low to avoid substantial lengthening of the weak sarcomeres and development of sarcomere length inhomogeneity. I-band titin could exert an efficient role in preventing this problem only if its stiffness got much larger during contraction.

A quantitative description of the I-band titin spring contribution *in situ* in the active half-sarcomere is hampered by the presence, in parallel, of the array of myosin motors with a stiffness that is more than one order of magnitude larger than titin stiffness (Fusi et al., 2014; Powers et al., 2020). In order to investigate titin mechanical properties in the different functional states of the half-sarcomere, we use Para-Nitro-Blebbistatin (PNB,(Kovacs et al., 2004; Limouze et al., 2004; Varkuti et al., 2016), to inhibit actin-myosin interaction even during tetanic stimulation of a frog muscle fibre. The action of blebbistatin in previous experiments was characterised by some residual motor activation in mammalian muscle (Fusi et al., 2016; Iwamoto, 2018; Ma et al., 2018), therefore we preliminarily defined the PNB concentration (20 μM) able to completely suppress the mechanical response and preserve the resting structure of thick filament of an intact frog muscle fibre upon tetanic stimulation and stretch. Under these conditions we exploited our fast sarcomere-level mechanics to determine the titin responses to stepwise changes in force imposed on the resting and stimulated fibre in the physiological SL range. We discovered that upon stimulation titin switches from the OFF-state characterised by a large, SL-dependent, extensibility, to the ON-state in which it acquires rectifying properties, opposing stretching with an effective stiffness of 4 pN nm^−1^ per half-thick filament and independent of SL, while preserving the ability of the half-sarcomere to shorten at velocity more than twice the unloaded shortening velocity exhibited by the muscle fibre in active contraction. The OFF-ON transition kinetics revealed to be strictly related to the Ca^2+^-dependent structural changes in the thin filament, indicating that the switching mechanism depends on titin binding to the thin filament that reduces the length of the I-band titin spring to that of the one third distal segment of the PEVK domain.

## RESULTS

### The myosin inhibitor PNB suppresses thick filament activation and motor attachment during tetanic stimulation and forced lengthening

Addition of 20 μM PNB to Ringer solution induces, within 30-40 min, the complete suppression of the force generated by single muscle fibres under tetanic stimulation (Figure 2A, blue in normal Ringer solution, orange in the presence of PNB). The structural correlate of the action of PNB at the level of the thick filament and myosin motors was tested by collecting 2D X-ray diffraction patterns at the ID02 beamline of the European Synchrotron (Grenoble, France). For the purpose of this work, low angle X-ray patterns collected at SL 2.7 μm (see STAR Methods) in normal Ringer solution (control, Figures 2B–2H, black at rest, blue at the plateau of the isometric tetanus, *T*_0_) were compared with those in the presence of PNB (brown at rest, orange during tetanic stimulation). Tetanic stimulation in control produces: (*i*) increase of the intensity ratio of the low angle equatorial reflections (*I*_1,1_/*I*_1,0_) from the resting value of 0.16 to the *T*_0_ value of 0.58 (Figures 2B and 2E), indicating movement of the myosin motors from the surface of the thick filament toward the thin filament (*ii*) reduction of the intensity of the ML1 layer line (*I*_ML1_) to ¼ (Figures 2C and 2F), indicating that the motors have lost the helical order on the surface of the thick filament and are either disordered or attached to actin; (*iii*) reduction of the intensity of the meridional reflections M1-M6, indexing on an axial periodicity of ~ 43 nm (the myosin-based reflections, Figure 2D), with the exception of the M3, associated with the myosin heads, which remains strong; (*iv*) change of M3 fine structure (arising from X-ray interference between the two arrays of myosin motors in each thick filament (Linari et al., 2000)), from a main peak at 14.35 nm and small satellite peaks on either side (black) to two peaks of similar intensities (blue). The ratio of the high angle peak intensity over M3 total intensity (*H*_M3_, Figure 2G) is 0.15 at rest (black) and becomes 0.50 at *T*_0_ (blue), indicating that myosin motors have moved from their resting configuration in which they lie on the surface of the thick filament tilted back in the OFF or IHM state (inset in Figure 1C, (Reconditi et al., 2011; Woodhead et al., 2005)) to either disordered or attached to actin tilted near the perpendicular to the filament axis (Brunello et al., 2007; Dobbie et al., 1998; Piazzesi et al., 2007; Piazzesi et al., 2002; Reconditi *et al*., 2011); (v) increase of the spacing of M6 reflection (*S*_M6_, Figures 2D and 2H) related to a periodicity in the thick filament backbone (Huxley et al., 2006) from 7.176 nm (black) to 7.271 nm (blue), 5 times larger than those expected on the basis of the elastic extensibility of the filament (Brunello et al., 2014; Reconditi et al., 2004) and correlated with slower stress-induced structural changes in the thick filament (Linari et al., 2015).

**Figure 2.**
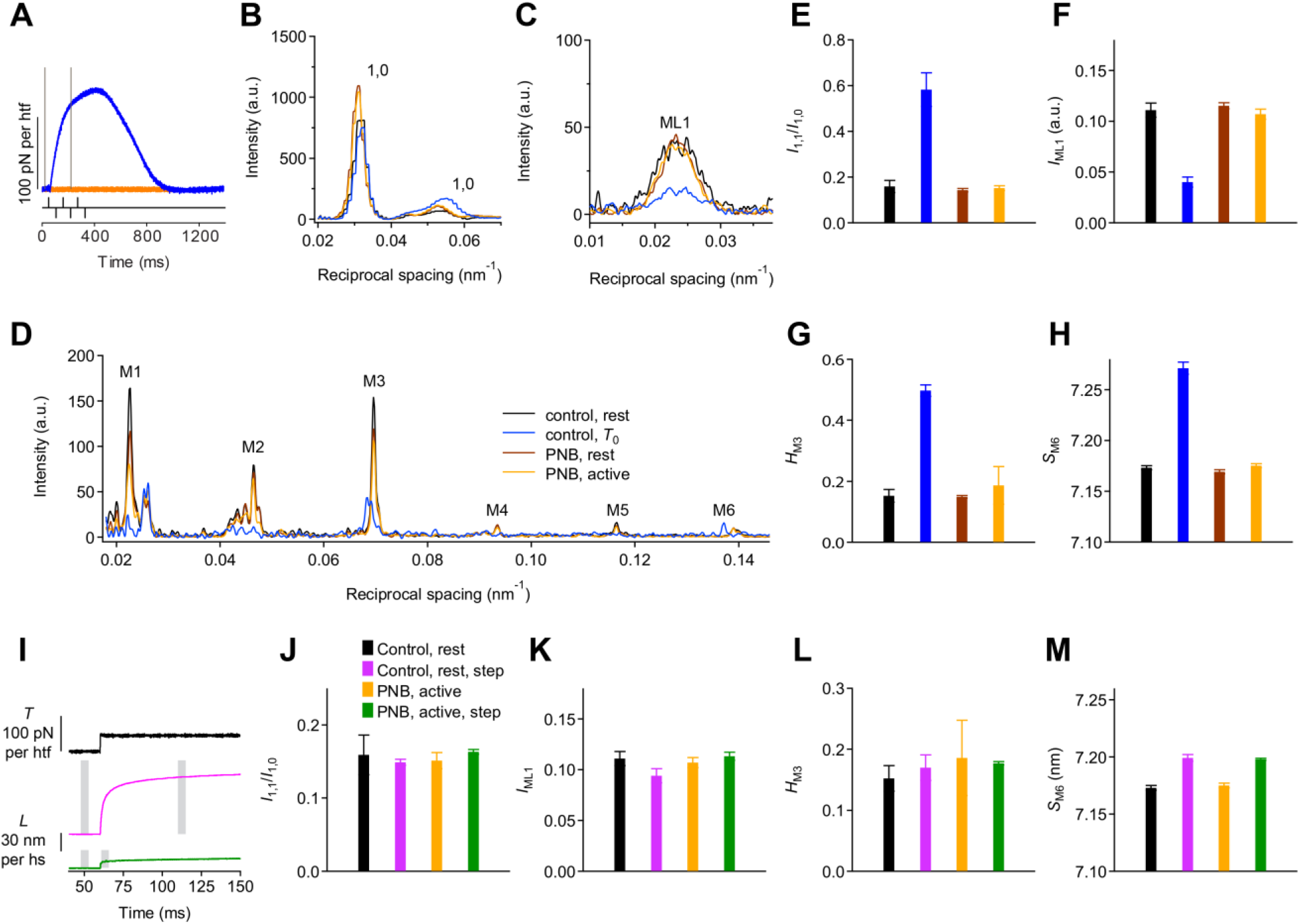
The force inhibitor PNB suppresses thick filament activation and motor attachment during tetanic stimulation and forced lengthening. **A.** Force response to tetanic stimulation (lower black trace) at 2.7 μm SL, either in physiological solution (control, blue trace) or after the addition of 20 μM PNB (orange). Grey bars: X-ray exposure time windows. **B-D**. X-ray diffraction intensity profiles at rest (black, control; brown, PNB) and during tetanic stimulation (blue, control; orange, PNB). Equatorial 1,0 and 1,1 reflections (B), first myosin layer line (ML1, C) and myosin-based meridional reflections, (M1-M6, D). **E**. Ratio of the intensity of the 1,1 reflection over the intensity of 1,0 reflection (*I*_1,1_/*I*_1,0_). **F**. Intensity of ML1 reflection (*I*_ML1_). **G**. Intensity ratio of the high angle peak of the M3 reflection over the total M3 (*H*_M3_). **H**. Spacing of the M6 reflection (*S*_M6_). The colours in the E-H histograms identify the same conditions as in B-D. **I-M.** Lengthening response to a force step imposed on the fibre either at rest in control (magenta) or at 60 ms following the first stimulus of tetanic stimulation in the presence of PNB (green) (I) and corresponding changes in the X-ray signals (J: *I*_1,1_/*I*_1,0_, K: *I*_ML1_, L: *H*_M3_, M: *S*_M6_). Black and orange histograms, same data and colour code as in B-H; magenta and green, X-ray signals in response to the step at rest in control and during stimulation in PNB, respectively. Temperature 4 °C. Error bars are SEM. Data in panels E-H and J-M are from 8 bundles, 4 for controls and 4 in PNB (*n*= 6-18).

The perfusion of PNB Ringer solution does not affect the X-ray signals at rest (compare brown and black in Figures 2B–2H) of either equatorial reflections (B and E), or ML1 (C and F) or meridional reflections (D, G and H). *t* test for the difference gives always P> 0.13). In PNB Ringer solution, upon tetanic stimulation all the X-ray signals (Figures 2B–2H, orange) maintain the values characteristic of the resting state (brown) also at the time when in control the force attains the *T*_0_ value (Figure 2A, blue). Thus, PNB preserves the OFF state of the thick filament also during the rise of internal [Ca^2+^] that in control elicits *T*_0_.

The effect of stretch on myosin motors in the absence/presence of PNB was tested by comparing the X-ray signals during the lengthening in response to a force step of ~0.25 *T*_0,*c*_ (the isometric tetanic force developed in the control solution, at 4 °C and 2.15 μm SL) either at rest in control (Figure 2I, magenta) or upon tetanic stimulation in the presence of PNB (green). As shown in Figures 2J–2M, either at rest in control (black before step, magenta after step) or during stimulation with PNB (orange before step, green after step) no significant changes in the X-ray signals marking the OFF state of the myosin motors accompany the response to the force step, except a ~0.35 % increase of the *S*_M6_ both at rest (Figure 2M, magenta versus black) or during tetanic stimulation (green versus orange).

This increase is ~5 times larger than that expected from the elastic extensibility of the filament and underpins the stress-induced structural transition occurring in the thick filament in the milliseconds time scale (Ma *et al*., 2018; Reconditi et al., 2019; Reconditi *et al*., 2004), confirming that this transition is Ca^2+^ independent (Ma *et al*., 2018). The small reduction of *I*_ML1_ accompanying the response to the force step at rest (Figure 2K, magenta versus black, −20%, P= 0.023) is not attributable to the force change itself, but rather to the extent of lengthening, as observed when a muscle at rest is brought to SL higher than 2.7 μm (see Figure 6A in (Reconditi *et al*., 2014)). In conclusion the X-ray analysis gives evidence that in the fibre activated in PNB as in the fibre at rest in control, the force step does not produce any alteration of the OFF state of myosin motors.

### During tetanic stimulation the I-band titin extensibility reduces by one order of magnitude and becomes independent of SL

The lengthening responses of ~500 sarcomeres, selected near the force transducer end of the fibre, to force steps of amplitude Δ*T*= 0.12 *T*_0,c_ (Figure 3A), 0.2 *T*_0,c_ (3B) and 0.4 *T*_0,c_ (3C) were recorded at different starting sarcomere lengths (range 2.3 – 3.0 μm) both at rest (dark colours) and during tetanic stimulation (light colours) at 4 °C. At rest, even for the smallest step, the lengthening response at 2.3 μm SL (dark grey in A) is so large that saturates the range of movement allowed by the loudspeaker-motor (± 600 μm, see STAR Methods). For the 0.12 *T*_0,c_ step (A) the lengthening response could be almost completely recorded at SL= 2.5 μm (blue) and fully recorded at SL 2.7 μm (dark green) and 3 μm (brown), showing that it is made by a fast, roughly exponential component complete within ~50 ms followed by a slow component at constant velocity, *V*_3_, ≤ 20 nm s^−1^ per half-sarcomere (hs). The amplitude of the fast component in response to a force step progressively decreases at larger SL. A quantitative estimate of the fast component amplitude (*L*_2_) is obtained as the ordinate intercept of the tangent to the later linear phase back-extrapolated to the time of the step (see STAR Methods). *L*_2_ and *V*_3_ for the 0.2 *T*_0,c_ reduce with the increase in SL from 2.7 to 3.0 μm, being 92 ± 5 nm and 22 ± 2 nm s^−1^ per hs respectively at 2.7 μm and 46 ± 4 nm and 11 ± 1 nm s^−1^ per hs respectively at 3.0 μm (see also Table S1, 4 °C, Rest). In the 22 fibres used in this work *T*_0,c_ is 144 ± 18 kPa (mean ± SD). From the lattice geometry of the frog muscle fibre *T*_0,c_ per half thick filament (htf) can be calculated and is 245 ± 32 pN (mean ± SD). Thus a force step of 0.2 *T*_0,c_ corresponds to ~50 pN per htf. The low lengthening velocity of the later phase (22.2 ±2.2 and 11.3 ±1.2 0 nm s^−1^ per hs at SL 2.7 and 3 μm respectively, see also Table S1, 4°C, Rest) indicates a viscous-like response with a drag coefficient as high as (Δ*T*/*V*3 =) 2.5-5 pN s nm^−1^. The effective resistance to the increase in load is therefore defined by the in series quasi-instantaneous stiffness *e*_2_ (= Δ*T*/*L*_2_). *e*_2_ is 1.10 ±0.09 pN nm^−1^ per htf at 3.0 μm SL and reduces to ½ (0.54 ±0.04 pN nm^−1^) at 2.7 μm SL (Figure 3D open circles, see also Table S1). At 2.5 μm SL *e*_2_ can be only deduced from the response to the 0.12 *T*_0,c_ step (Figure 3A, blue) and is (30/120 =) 0.25 pN/nm per htf (Figure 3D, filled circle).

**Figure 3.**
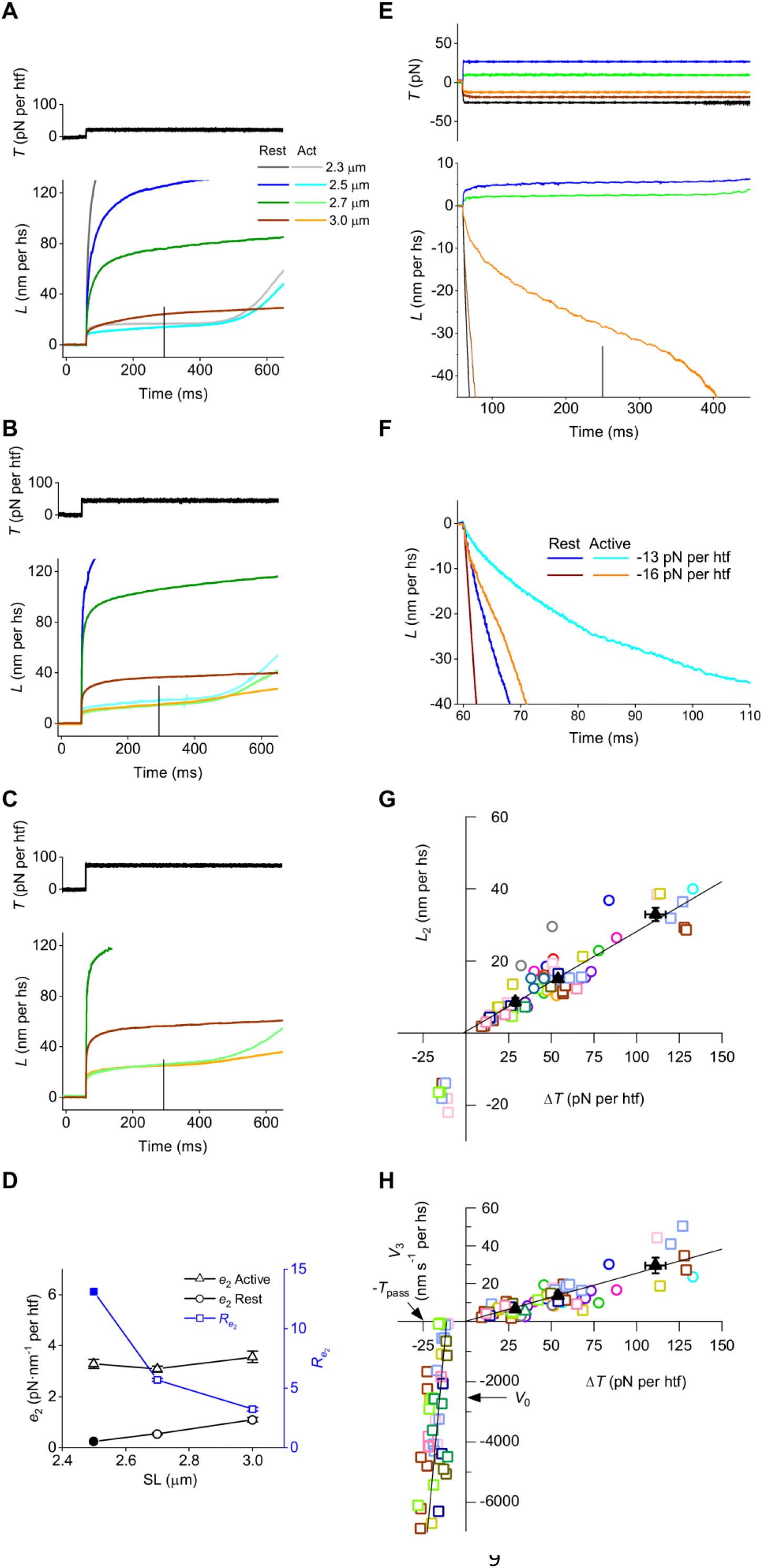
During tetanic stimulation the I-band titin becomes a mechanical rectifier resisting to pulling forces and shortening freely under restoring forces. **A-C.** Lengthening responses (lower superimposed traces in each panel) to positive force steps (upper traces, black) of 0.12 *T*_0,c_ (A), 0.20 *T*_0,c_ (B) and 0.40 *T*_0,c_ (C) imposed in the presence of PNB either at rest (dark colours) or during stimulation (light colours) at different SL (see also Table S1): dark and light grey, 2.3 μm; blue and cyan, 2.5 μm; dark and light green, 2.7 μm; brown and orange, 3.0 μm. Stimulation starts at 0 ms, ends at 290 ms (vertical line); force step is imposed at 60 ms. **D.** *e*2 (left ordinate) against SL at rest (circles) and during stimulation (triangles) estimated with a force step of ~50 pN (open symbols, from Table S1); the filled circle is *e*_2_ with ~30 pN steps imposed at 2.5 μm SL. Blue squares (right blue ordinate): ratio between *e*2 in active and resting fibres (*R*_e2_). **E**. Superimposed length changes (lower traces) in response to force steps (upper traces) of different sizes and directions (as indicated by figures close to traces) imposed during tetanic stimulation at 3 μm SL (see also Tables S2 and S3). **F**. Comparison of shortening responses to negative force steps imposed on a fibre either at rest (dark colours) or during stimulation (light colours) at 3 μm SL (−13 pN per htf, blue and cyan; −16 pN per htf, brown and orange). **G.** *L*_2_ - Δ*T* relation at 3 μm SL. Different colours identify different fibres on which either only positive steps (circles) or both positive and negative steps (squares) were imposed. Black filled triangles: mean ± SEM for positive steps >20 pN grouped in three classes of forces (from Table S2). The line is the linear fit on pooled data (slope 0.277 ± 0.016 nm pN^−1^, intercept 0.45 ± 0.97 nm per hs). **H.** *V*_3_ - Δ*T* relation at 3 μm SL. Symbols with the same colour code as in G. The line in the right upper quadrant is the linear fit to pooled data (slope 0.254 ± 0.021 nm s^−1^ pN^−1^, intercept 0.11 ± 1.25 nm s^−1^ per hs). The line in the left lower quadrant (slope 624 ± 135 nm s^−1^ pN^−1^, intercept 7042 ± 1643 nm s^−1^ per hs) is obtained as the average of the linear fits to data from individual fibres (see STAR Methods). The arrows indicate the static passive force (*T*_pass_) and the unloaded shortening velocity in control contractions (*V*_0_). All data collected at 4 °C.

The response to the force step during tetanic stimulation (light colours in Figures 3A–3C) is characterised by an *L*_2_ that is much smaller and faster (half time within 500 μs) than that at rest. The later steady lengthening occurs at a slow speed (*V*_3_ ~12 nm s^−1^ per hs, see also Table S1) until, following the end of stimulation (vertical bar at 290 ms), it undergoes a sharp transition to a faster lengthening. The time elapsed from the end of stimulation to this transition (*t*_OFF_, see STAR Methods and also Figure S2) is a clear indication of the delay with which the extensibility of the sarcomere at rest is resumed. Most strikingly, *L*_2_ for a given force step in the active fibre is almost independent of SL (Figures 3B–3C, light colours, see also Table S1, 4 °C, Active; P of the differences is always >0.28). The corresponding *e*_2_ ranges from 3.1 pN nm^−1^ per htf to 3.6 pN nm^−1^ per htf, but the difference is not statistically significant (P> 0.05). Also *V*_3_ (~11.4 – 12.7 nm s^−1^ per hs) does not significantly depend on SL (P of the differences is always >0.08), underlying a drag coefficient of (50 pN / 12 nm s^−1^ =) 4 pN s nm^−1^.

The comparative analysis of the dependence of the lengthening response on the step size in resting and active fibres, possible only at 3 μm SL, shows that (*i*) in both cases *L*_2_ increases with the step size, from 31.4 (with 30 pN) to 72 nm per hs (with 94 pN, see also Table S2), so that *e*_2_ is not significantly affected (P always > 0.20); (*ii*) *e*_2_ is three times smaller at rest (1 - 1.4 pN nm^−1^) than during activation (3.4 - 3.8 pN nm^−1^). Thus, the extensibility of the hs at rest, which under our experimental conditions represents the extensibility of the I-band titin, reduces under tetanic stimulation by a factor that is 3 at 3 μm SL, more than 5 at 2.7 μm and likely more than 10 at 2.5 μm (Figure 3D, blue squares). At 2.3 μm SL, at which the saturation of movement of the motor prevents any quantitative evaluation, the factor is likely more than 20 (compare dark and light grey traces in Figure 3A).

At rest *e*_2_ is smaller at shorter SL because of the larger extension required to straighten the poly-Ig domain to get its contour length *L*_c_, ~3.0 μm (Linke *et al*., 1998b; Trombitas *et al*., 1998), Figure 1C-D, see also Note S1). Under this condition, the lengthening in response to a force step applied at 3.0 μm will mainly elicit the elastic strain of the PEVK domain.

We conclude that stimulation switches the I-band stiffness from an OFF-state characterised by a large SL-dependent extensibility to an ON-state characterised by a high, SL independent resistance to stretch, made of an elastic component with stiffness >3 pN nm^−1^ per htf in series with a viscous component with a drag coefficient ~4 pN s nm^−1^. The underlying mechanism must rely on an activation-dependent process that, even at SL ≪3 μm excludes the contribution of the compliant poly-Ig domain. Supposedly, this process consists in the formation of a link between a point in the titin distal to the poly-Ig domain and the nearest actin monomer in the thin filament, so that the compliant poly-Ig domain is substituted with the much stiffer thin filament segment.

### Titin in the ON state acts as a mechanical rectifier

Whatever the mechanism, the transition in the I-band titin responsible for the rise in the resistance to stretch should not imply any significant resistance to muscle shortening, as the maximum velocity of shortening of active muscle (the velocity of unloaded shortening, *V*_0_) is constant independent of the number of motors available for the interaction with actin (Edman et al., 1979; Fusi et al., 2017; Linari *et al*., 2015). The question is investigated here by recording the response to stepwise drops in force imposed at 3 μm SL, at which the steady passive force of ~0.1 *T*_0,c_ (Figure 1B) is enough to make the measure feasible. In the experiment of Figure 3E, with the smallest force step (negative step, orange; positive step, green) the shortening transient maintains the multiphase aspect of the lengthening transient, even if phase 2 is much larger and slower evolving to a steady velocity (*V*_3_), ~-70 nm s^−1^ per hs, also larger. Following the end of stimulation (vertical bar at 300 ms) the shortening velocity increases with a delay (*t*_OFF_) similar to that determined in the lengthening transient (Figures 3A–3C). Increasing the size of the step by a few pN per htf, while the lengthening response maintains the same features (blue), the shortening responses (brown and black) become monotonic as *V*_3_ increases abruptly merging with phase 2.

*V*_3_ elicited by a negative force step imposed at rest at 3 μm SL is 2-3 times larger than in the stimulated fibre (Figure 3F, dark colours rest, light colours active). Moreover, even for the smallest step (−13 pN) the response of the resting fibre (blue) does not show any sign of the *L*_2_ component of the active response (cyan). Data from 8 fibres for the same range of steps (−0.035 - −0.065 *T*_0,c_) give a *V*_3_ of the resting fibre (−6433 ± 1093 nm s^−1^ per hs) more than three times larger than that of the active fibre (−1780 + 443 nm s^−1^ per hs, see also Table S3). Thus, activation induces a drag coefficient for shortening that, even if very little, is three times larger than in the resting fibre.

The dependence on the step size and direction of *L*_2_ and *V*_3_ in the active fibre at 4 °C are shown in Figures 3G and 3H respectively (data-points for steps larger than 20 pN are those used for the averages, black filled triangles). The linear regression to pooled *L*_2_ - Δ*T* data for positive steps has a slope of 0.277 ± 0.016 nm pN^−1^. Its reciprocal (an estimate of *e*_2_) is 3.61 ± 0.21 pN nm^−1^; the ordinate intercept, 0.45 ± 0.95 nm is not significantly different from zero, confirming the direct proportionality between *L*_2_ and Δ*T*. *L*_2_ for negative steps, which can only be recorded within a small range of forces (−13.16 ± 0.80 pN per htf), is −16.96 ± 1.07 nm per hs, underpinning an *e*_2_ of 0.8 pN/nm, more than four times smaller than that for positive steps. The *V*_3_ - Δ*T* relations for either negative or positive steps (Figure 3H, from the same fibres as in Figure 3G) appear quite linear (see STAR Methods for the different approaches used to estimate the slopes of the relations). The slopes, that estimate the fluidity of the system (the reciprocal of the drag), are 0.254 ± 0.021 nm s^−1^ pN^−1^ and 624 nm s^−1^ pN^−1^ for positive and negative steps respectively. The corresponding drag coefficients are 3.94 pN s nm^−1^ and 1.60·10^−3^ pN s nm^−1^ respectively. A striking feature that emerges from the analysis of the response of the active fibre to a negative force step is that, in the absence of myosin motors, the restoring force exerted by titin at 3 μm SL (−*T*_pass_ = −24.94 ± 1.52 pN per htf (mean ± SD), the abscissa value indicated by the arrow in Figure 3H), accounts for an unloaded shortening velocity that is more than twice the unloaded shortening velocity accounted for by myosin motors during active contraction (*V*_0_ = −2530 ± 90 nm s^−1^ per hs in these experiments at 4 °C, the ordinate value indicated by the arrow). Notably, the contribution of titin in the ON state explains the finding (Edman *et al*., 1979) that, at SL >2.8 μm, the unloaded shortening velocity of contracting muscle increases sharply with SL above the value of *V*_0_ exhibited in the SL range 1.8-2.7 μm.

The slope change across zero force step indicates that the drag opposed by titin during shortening is more than three orders of magnitude smaller than during lengthening, underpinning the definition of titin in the ON state as a mechanical rectifier, which opposes to half-sarcomere lengthening with a high resistance made by an elastic element with stiffness 3.6 pN nm^−1^ in series with a viscous element with a drag coefficient 4 pN s nm^−1^, while it allows high speed shortening as the drag coefficient becomes ~2500 times smaller. The intercept on the force axis calculated from the fit for negative steps (see STAR Methods) is −11.20 ± 0.88 pN. This, together with the presence of *L*_2_ for the smallest negative step (Figures 3E, orange, and 3G) and the ca. threefold decrease in *V*_3_ with respect to the resting value, gives further support to the idea that in the ON state the I-band titin links to the thin filament, provided that under negative stress the link becomes highly dynamic.

### Dependence on temperature of the ON-OFF transition kinetics

Repeating the 0.2 *T*_0,c_ force step protocol at 14 °C (Figure 4A) shows that temperature has no evident effects on the lengthening transient either at rest (4 °C, blue and 14 °C, brown) or during stimulation (4 °C, cyan and 14 °C, orange). As at 4 °C, *L*_2_ and *V*_3_ in the resting fibre decrease with SL and in the active fibre do not change significantly with SL (Table S1, P always > 0.02). Two features of the lengthening transient of the active fibre are affected by temperature in a similar way at any SL: (*i*) *V*_3_ increases significantly (Table S1, P always <0.01): the average from the three SL’s is 12.1 ± 0.6 nm s^−1^ per hs and 19.1 ± 0.7 nm s^−1^ per hs at 4 °C and 14 °C respectively; (*ii*) *t*_OFF_ reduces to less than ½: the average from the three SL’s is 191± 13 ms and 70± 3 ms at 4 and 14 °C respectively (see also Table S4).

**Figure 4.**
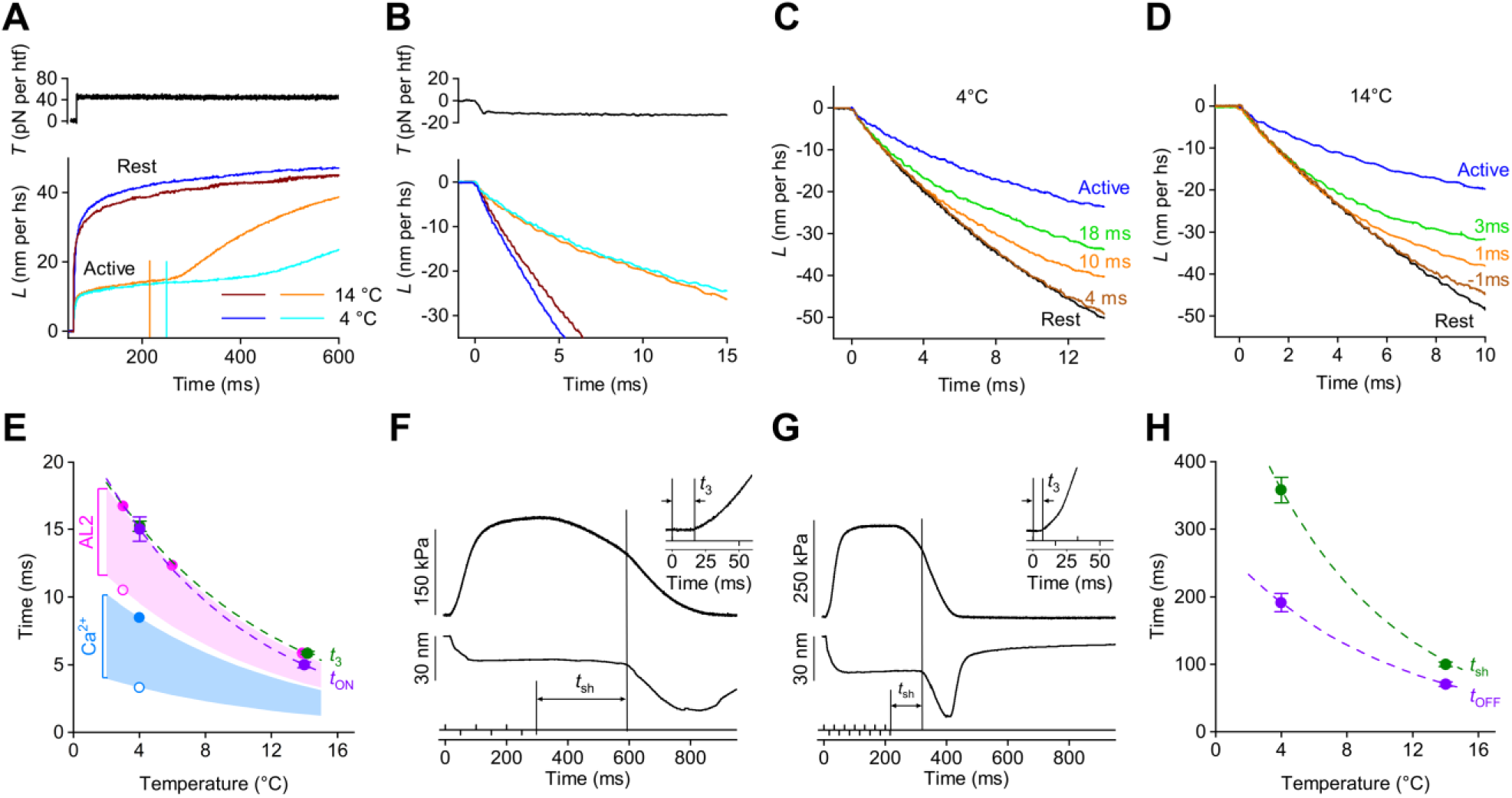
The ON-OFF transition kinetics of the I-band titin and its temperature dependence are accounted for by thin filament activation. **A.** Lengthening responses (lower traces) following a positive force step (50 pN, upper trace) imposed either at rest (dark colours) or during stimulation (light colours): blue and cyan 4 °C, brown and orange 14 °C. Stimulus start, 0 ms; end, 250 ms (4 °C, cyan vertical line) and 216 ms (14 °C, orange vertical line). **B.** Shortening responses (lower traces) following a negative force step (−12 pN, upper trace) under conditions defined by the same colour code as in A. **C**. Superimposition of shortening responses at 4 °C to a force step of −12 pN imposed at rest (black) and at different times (Δ*t*) from the first stimulus: 4 ms, brown; 10 ms orange; 18 ms, green and 60 ms (active), blue. **D**. Superimposition of shortening responses at 14 °C to a force step of −12 pN imposed at rest (black), at 1 ms before the first stimulus (brown) and at different times from the first stimulus:1 ms, orange; 3 ms, green and 60 ms (active), blue. SL in panels A-D, 3.0 μm. **E.** Temperature dependence of the ON transitions of the relevant parameters (identified by the colour, see also Table S4), with *t*_ON_ defined as the time from the foot of the action potential or the first stimulus to the onset of the parameter change: Blue, Ca^2+^ transient: open circle, *t*_ON_; filled circle, *t*p (time to Ca^2+^ peak); shaded area, temperature dependence of the Ca^2+^ rise. Magenta, AL2 increase: open circles *t*_ON_; filled circle, *t*1/2; shaded area, temperature dependence of AL2 increase. Violet circles and dashed line, latency of titin switching ON (*t*_ON_) and its temperature dependence. Green circles and green dashed line, latency of force development (*t*_3_) and its temperature dependence. **F** and **G.** Contraction-relaxation cycle of a tetanus in control at 2.15 μm SL at 4 °C (F) and 14 °C (G). Upper trace, force; middle trace, half-sarcomere length change; lower trace, stimuli. The long vertical line marks the start of chaotic relaxation at a time *t*_sh_ after the last stimulus (short vertical line). Inset: expanded force trace of the corresponding panel to show *t*_3_, the latency for force rise. **H**. Temperature dependence of the OFF transitions (see also Table S4): violet circles and dashed line: *t*_OFF_ and its temperature dependence. Green circles and dashed line, *t*_sh_ and its temperature dependence.

As for the lengthening response, also the shortening response is scarcely affected by 10 °C increase in temperature, either at rest (Figure 4B, 4 °C blue, 14 °C brown) or during stimulation (4 °C cyan, and 14 °C orange). Within the same range of negative step sizes as at 4 °C, at 14 °C *V*_3_ of the resting fibre (−7078 ± 683 nm s^−1^ per hs) is more than three times larger than that estimated in the active fibre (−2009 ± 701 nm s^−1^ per hs,) and in either condition is not significantly different from *V*_3_ at 4 °C (, P always >0.6, see also Table S3). Thus a 10 °C increase in temperature does not significantly affect the viscous component of shortening either at rest or (unlike lengthening) during stimulation.

The lengthening response to a positive force step is able to record the ON-OFF transition of the I- band titin spring following the end of stimulation through the parameter *tOFF*, demonstrating that the kinetics of the switching OFF is temperature dependent (Figure 4A, cyan versus orange, see also Tables S1, S2 and S4) with a Q_10_ of 2.70.

The lengthening transient, however, cannot *per se* have the time resolution to describe the OFF-ON transition at the start of stimulation, because the fast phase of the lengthening transient takes a few tens of milliseconds to attain *L*_2_, a time that might be comparable to that taken by the Ca^2+^-dependent activation. Instead, the resting-active difference of the shortening velocity in response to a negative force step imposed at 3 μm SL emerges within few hundred microseconds after the step as shown in Figure 3F by the superimposition of corresponding resting (dark colours) and active (light colours) responses. This feature is used as a tool to estimate the time after the start of stimulation at which the OFF-ON transition starts (*t*_ON_), by superimposing the shortening responses obtained with different delays between the start of stimulation and the imposed force step (Δ*t*) (Figure 4C). Within the 10 ms of shortening time window, the switching ON appears as a deviation of the trace from that of the resting fibre (black) toward that of the active fibre (blue). The time at which the deviation occurs, which decreases by increasing Δ*t*, is more precisely estimated by the difference trace (*DL*) between the resting response and the responses obtained with different Δ*t* (see STAR Methods). *t*_ON_, calculated by adding Δ*t* and the time after the step for the upper deviation of *DL* from the baseline, is 15.03 ± 0.89 ms at 4 °C (*n* = 17 from four fibres). At 14 °C (Figure 4D), for the deviation to occur within the 10 ms time window, Δ*t* must be ~10 ms shorter than at 4 °C. *t*_ON_ at 14 °C is 4.98± 0.22 (*n* =22, from the same four fibres plus two more fibres, see also Table S4).

### The kinetics of OFF-ON transition of the I-band titin is accounted for by Ca ^2+^-dependent thin filament activation

The hypothesis that the OFF-ON switch in the I-band titin spring is induced by the rise of [Ca^2+^]_i_ can be tested in a comparative analysis of the time courses of the OFF-ON titin transition and of the Ca^2+^-related signals accompanying the contraction-relaxation of the frog muscle fibre (Figures 4E–4H).

In frog muscle fibres at 4 °C [Ca^2+^]_i_ starts to rise at 3.3 ms (*t*_ON_, blue open circle in Figure 4E) after the foot of the action potential and attains its peak at 8.5 ms (*t*_1/2_, blue filled circle) (Caputo et al., 1994; Sun et al., 1996). By taking into account a Q_10_ of 2.5 for the Ca^2+^ transient (Eusebi and Miledi, 1983; Miledi et al., 1982) it is possible to define the time course of calcium transient in the range of temperatures of the present experiments (4-14 °C, blue shaded area in Figure 4E).

Ca^2+^ binding to Troponin (Tn) triggers the azimuthal movement of Tropomyosin (Tm) around the thin filament surface that makes the actin sites available for binding of the myosin motors (Gordon et al., 2000). The Tm movement has been recorded in frog muscle by the increase in the intensity of the 2^nd^ actin layer-line (AL2), sensitive to the position of Tm in the thin filament (Kress et al., 1986). At 3 °C, the square root of AL2 intensity, which measures the time course of the structural changes, starts to rise at 10.5 ms from the first action potential (*t*_ON_, magenta open circle at 3 ms in Figure 4E) and attains its maximum with a half-time of 16.7 ms (*t*_1/2_, magenta filled circle). The half-time reduces increasing the temperature (magenta filled circles at 6 and 14 °C). The relative Q_10_ (2.63) is used to calculate how the latency for the Tm movement reduces with temperature and draw the magenta shaded area that defines the time course of Tm movement for the range of temperatures between 3 and 14 °C. Noteworthy, the start of Tm movement almost coincides with the time for the peak of the Ca^2+^ transient. How these signals relate to the switching ON of the I-band titin can be defined by reporting on the same graph (Figure 4E) the temperature dependence of *t*_ON_ (violet circles and dashed line, see also Table S4). We conclude that switching ON of titin, whatever the mechanism, starts when Tm movement is half-complete, in spite of the much faster rise in [Ca^2+^]_i_. If switching ON of titin upon activation is due to formation of the actin-titin link, it follows that titin binds an actin site only when it is released from the steric inhibition of Tm.

The same mechanism is known to operate for starting actin-myosin interaction. The time elapsed between the start of depolarization and the start of force rise in the isometric tetanic contraction (Figures 4F and 4G), defined as *t*_3_ in the inset according to (Mulieri, 1972), has a marked temperature dependence being 15.26 ± 0.38 at 4 °C and 5.87 ± 0.12 at 14 °C (see also Table S4). *t*_3_ and its temperature dependence (Figure 4E green circles and dashed line respectively) are associated to Tm movement just like titin *t*_ON_. Notably *t*_3_ coincides also with the time required to develop the maximum shortening velocity in unloaded conditions (*V*_0_), a property that requires only a few myosin motors available for interacting with actin and precedes the much slower load-dependent massive recruitment of the myosin motors from their resting state along the thick filament (Fusi *et al*., 2017; Huxley and Hanson, 1957; Linari *et al*., 2015).

The temporal relation between the return of titin to the resting state (OFF transition) and the mechanical relaxation is shown in Figure 4H. In a frog muscle fibre the decay of [Ca^2+^]_i_ to near resting values following the end of stimulation is much briefer than the mechanical relaxation (Caputo *et al*., 1994). A straightforward quantitative information on the ensuing movement of Tm back to the resting position in the thin filament is not available, because its time course during contraction is affected by the presence of actin-attached myosin motors (Kress *et al*., 1986; Matsuo and Yagi, 2008). In this respect, the use of PNB is the ideal tool to release the OFF transition in the thin filament structure from the influence of actin-attached motors. *t*OFF, our estimate of the time of titin switching OFF, reduces from 191 ± 13 ms at 4 °C to 70 ± 3 ms at 14 °C (Figure 4H, violet circlesand violet dashed line, calculated with the corresponding Q_10_, see also Table S4).

Relaxation of the isometric force following the end of tetanic stimulation (Figures 4F and 4G at 4 and 14 °C respectively) is much slower and accelerates only at a time after the last stimulus at which the tension shows a shoulder (*t*_sh_) (Huxley and Simmons, 1970). *t*_sh_ (marked by the vertical line in Figures 4F and 4G) is constant independent of SL and decreases with temperature going from 358 ± 19 ms at 4 °C to 100 ± 3 ms at 14 °C (see also Table S4). This slow phase of relaxation is characterised by almost no change in half-sarcomere length (Figures 4F and 4G, middle trace) and myofilaments maintaining their ON structure (Brunello et al., 2009), while the acceleration of the relaxation at *t*_sh_ is characterised by chaotic behaviour of sarcomere populations with some of them shortening and others giving (Brunello *et al*., 2009). *t*_sh_ (green circles and dashed line in Figure 4H) follows the switching OFF of I-band titin spring (violet), which is expected if the recovery of the resting I-band titin extensibility at any SL is the necessary condition for the transition to the chaotic and rapid phase of relaxation.

## DISCUSSION

### Switching ON of I-band titin is explained by binding of the proximal two-thirds of the PEVK domain to the actin filament

The extensibility of the I-band titin spring in the fibre at rest is incommensurably large at full filament overlap and progressively reduces with the increase in SL. This feature is the signature of the OFF-state of titin, in which the I-band titin elasticity is governed by the large extensibility of the poly-Ig domain until this portion attains its contour length, which putatively occurs at SL ~3 μm (Figure 1D, Figure S1). The high, SL independent, resistance to stretch (~3.6 pN nm^−1^ per htf) and the ability to shorten at a velocity more than twice the unloaded shortening velocity of the contracting sarcomere are the signature of the ON-state of titin. These properties entail the characteristics of a mechanical rectifier that opposes to positive, pulling forces (drag coefficient 3.9 pN s nm^−1^), while under negative pushing forces it shortens freely with a three order of magnitude smaller drag coefficient (1.6·10^−3^ pN s nm^−1^). The time courses of both the OFF-ON transition following the start of stimulation and ON-OFF return after the end of stimulation are temperature sensitive (Figures 4E and 4H). Instead, most of the mechanical features marking the OFF and ON states of the I-band titin do not depend on temperature (Figures 4A and 4B, see also Tables S1 and S2), with the exception of the velocity of lengthening (*V*_3_) following a positive force step imposed on the active fibre.

The finding that in the ON-state the I-band titin spring exhibits a resistance to stretch that is independent of SL and three times higher than that in the OFF-state at 3 μm SL poses fundamental constraints on the underlying mechanism. Upon thin filament activation a region in the I-band titin distal to the poly-Ig domain (Figure 5) binds to the nearby actin site and excludes the poly-Ig domain viscoelasticity (magenta box in Figure 5C) from contributing to the mechanical response, by substituting it with the much stiffer thin filament. In this way the stiffness in the ON-state becomes independent of the SL-dependent extensibility of the poly-Ig domain.

**Figure 5.**
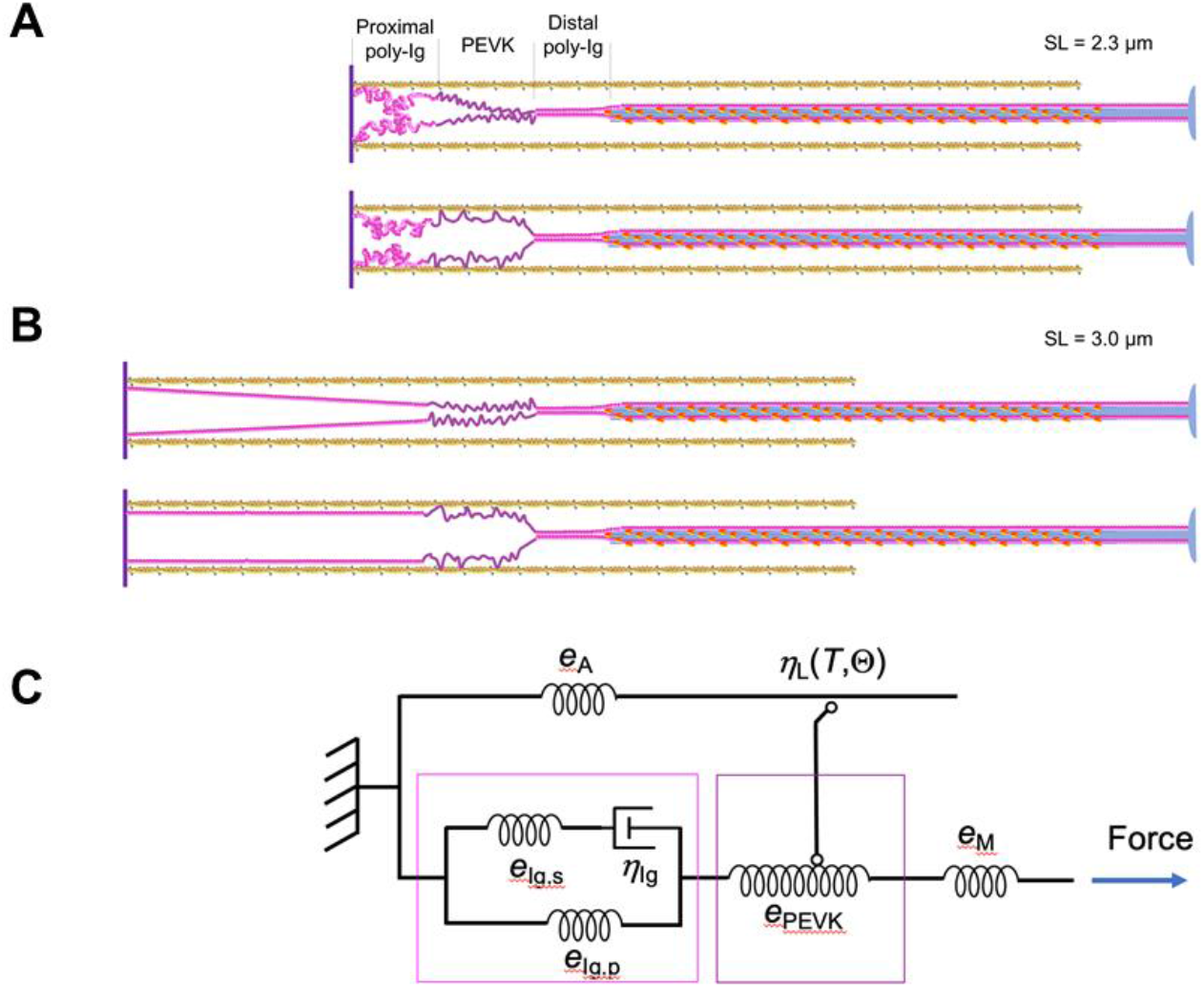
Overview of the OFF and ON states of titin and equivalent mechanical model. **A**. OFF (top) and ON (bottom) states at 2.3 μm SL. The proximal 2/3 of the PEVK domain (dark magenta) forms multiple links with the overlapping proximal segment of thin filament. **B**. OFF (top) and ON (bottom) states at 3.0 μm SL. The proximal 2/3 of the PEVK domain link the more distal overlapping segment of thin filament. Same colour code as in Figure 1C. **C**. Equivalent mechanical model of the half-sarcomere in the absence of attached myosin motors. *e*_A_ and *e*_M_, represent the stiffness of the thin and thick filaments respectively. The viscoelastic properties of the proximal poly Ig domain (magenta box) are synthetized by a Maxwell element, with viscosity ηIg and stiffness *e*_Ig,s_, in parallel with a spring with stiffness *e*_Ig,p_ responsible for the static restoring force at SL > 2.7 μm; *e*_PEVK_ represents the stiffness of the PEVK domain elasticity (dark magenta box). The switch represents the actin-titin link with the property of a stress (*T*) and temperature (Θ) dependent viscosity ηL (*T*, Θ).

Molecular evidence of the titin-actin interaction can be found in the literature regarding either the whole titin (Kellermayer and Granzier, 1996) or different titin fragments (Kulke et al., 2001; Nagy et al., 2004; Yamasaki et al., 2001). In general, the PEVK domain was found to have a specific affinity for actin even if its modulation by Ca^2+^ remained under question, apart a direct effect of Ca^2+^ on the PEVK bending rigidity that accounts for a 30% increase in sarcomere stiffness (Labeit et al., 2003). More recently a Ca^2+^-dependent affinity for the actin filament of a construct of the N2A domain, which in the skeletal muscle isoform links the poly-Ig domain and the PEVK domain, was reported (Dutta et al., 2018), but the Ca^2+^ induced changes in the rupture force of the N2A-actin bond were too small to explain the changes in the I-band titin stiffness found *in situ* in this work. The contradiction is explained if titin-actin interaction is controlled not by a direct Ca^2+^-binding to titin but by a Ca^2+^-dependent structural transition in the thin filament, proven here by the temporal relation between OFF-ON transition and Tm movement (Figure 4E), that is lost in the *in vitro* experiment.

This conclusion finds support in the recently published crystal structure of the N2A segment (Stronczek et al., 2021) that demonstrates the absence of specific Ca^2+^ binding sites in that domain. In the same paper no evidence for an N2A-actin interaction is found, attributing to the N2A region the sole function of signalosome. Eventually, an affinity of the N2A segment for actin has been found under the control of the muscle ankyrin repeat protein MARP1, which is overexpressed in the stressed muscle and specifically binds the N2A segment (van der Pijl and Ottenheijm, 2021; Zhou et al., 2021). These most recent contributions account for a modulation of the passive force-SL relation, responsible for the SL dependent increase in passive stiffness, but not for the activation-dependent change in titin stiffness reported here, which is independent of SL and, in the physiological range of SL, increases the resistance to stretch by one order of magnitude (Figure 3D). At 3 μm SL, at which the poly-Ig domain has attained its *L*_c_, I-band stiffness is threefold larger in the active fibre than at rest and depends on the stiffness of two essentially elastic elements in series (Figure 5): the actin filament proximal to the link (with stiffness *e*_A_) and the I-band titin distal to the link, mainly constituted by the PEVK domain (with stiffness *e*_PEVK_). With *e*_A_ >> *e*_PEVK_, the half-sarcomere stiffness is mainly attributable to PEVK domain and a threefold increase in *e*PEVK upon activation can be explained if titin attaches to actin within the proximal two-thirds of the PEVK domain, reducing the PEVK segment length to its distal third (Figure 5). This conclusion is strongly supported by the original *in vitro* experiments by Nagy et al (2004), showing that the PEVK sequence is constituted of two main motifs, the PPAK and the poly-E, of which the poly-E motif, which has the largest actin-affinity, is concentrated in the proximal two-thirds of the PEVK segment. This mechanism implies that the actin-titin interaction is made by multiple, in parallel, links, which is a compelling requirement considering the efficiency of the mechanism in resisting forces comparable to *T*_0,c_, the force exerted by the array of myosin motors during isometric contraction. Further viscous-like lengthening following a positive force step occurs at the low velocity *V*_3_ which, unlike the similar *V*_3_ of the resting fibre, depends on temperature (see also Table S1). Thus, in contrast to the poly-Ig based viscosity of the resting fibre (η_Ig_, Figure 5C), the viscous component (η_L_) of the response of the active fibre to the increase in force is mainly attributable to the titin-actin link dynamics, with a very low and temperature dependent detachment rate.

The stiffness of the I-band titin in phase 2 (*e*_2_) becomes so large upon activation to be comparable to that exhibited in the same time domain by the thick filament in series (*e*_M_, Figure 5C) (Ma *et al*., 2018; Reconditi *et al*., 2019; Reconditi *et al*., 2004). From the increase in *S*_M6_ following a 0.25 *T*_0,c_ step (0.36%, Figure 2M), the increase in the length of htf for a force of *T*_0_ is ((0.0036/0.25)*800 nm =) 11.52 nm. Taking *e*_2_ = 3.6 pN/nm per htf (Figure 3G) and *T*_0,c_ = 245 pN per htf, the extension of the I-band titin would be (245/3.6 =) 68.06 nm per *T*_0,c_. Correcting it for the contribution of thick filament brings the I-band titin extension to (68.06-11.52 =) 56.54 nm per *T*_0,c_ and the corrected *e*_2_ to 4.3 pN nm^−1^ per htf, a value 20% larger than the uncorrected one.

### How the dual state of titin bites into muscle contraction-relaxation

At SL up to 2.7 μm the stiffness of titin deduced by the passive force-SL relation (Figure 1B) is too low to prevent, during contraction at physiological SL, substantial lengthening of the weaker half-sarcomeres and development of SL inhomogeneity. Similarly, the increase by a few ten percent of the steepness of the passive force-SL relation by various modulators such as Ca^2+^ (Labeit *et al*., 2003) or PEVK phosphorylation (Hidalgo et al., 2009) cannot suit the task.

To quantify how efficiently the ON state of titin defined in this work plays against the intrinsic instability of serially linked half-sarcomeres, let’s assume that in a muscle fibre stimulated to develop an isometric tetanus at 2.5 μm SL a weak half-sarcomere undergoes a stress of 0.2 *T*_0,c_ (or 50 pN per htf) for the exceeding force of a strong half-sarcomere. Within the first 50 ms of contraction, the strain of the switched ON I-band titin spring in the weak half-sarcomere will be limited to (Δ*T*/*e*_2_= 50 pN/4.3 pN nm^−1^) = 11.6 nm. In a tetanus of 500 ms duration, further strain would be limited by the drag coefficient to (Δ*T*· *t* /η_L_ =50 pN · 0.5 s /3.9 pN s nm^−1^ =) 6.4 nm for a total strain of 18 nm. The average SL of the weak sarcomeres would have risen from 2.5 to 2.536 μm, that is by 1.4%. A semi-quantitative evaluation of the resistance to stretch at 2.3 μm SL (grey trace in the left column of Figure 3) indicates that also at full filament overlap the I-band titin in the ON-state responds under stress with a strain comparable to that reported for 2.5 μm SL and above.

Two other relevant issues emerge when the active titin properties discovered here are reflected on the physiology of muscle. The first derives from the finding that titin in the ON-state behaves as a mechanical rectifier resisting to a change in force with a drag coefficient that for negative force steps is three orders of magnitude smaller than for positive steps (Figure 3H). Accordingly, a drop to near zero of the restoring force present at 3 μm SL induces a *V*_3_ of ~6000 nm s^−1^ per hs, almost three times larger than the unloaded shortening velocity, *V*_0_, accounted for by the myosin motors in active contraction. Thus, titin does not exert any resistance against unloaded shortening velocity, which, in fact, is independent of number of motors and only depends on detachment kinetics of myosin motors (Edman *et al*., 1979; Fusi *et al*., 2017; Hanson and Huxley, 1957).

The second issue emerges in relation to the acceleration of the relaxation in the isometric tetanus. During the isometric phase of relaxation before the tension shoulder (Figures 4F and 4G) the [Ca^2+^]_i_ has already dropped near resting values (Caputo *et al*., 1994), while myosin motors detach very slowly as estimated by the slow decay of force and stiffness (Brunello *et al*., 2009). This can be explained by considering that motor detachment is strain dependent, increasing with the reduction of the load and, in this phase, attached motors are on average more strained than during isometric contraction as a consequence of the recoil of the series tendon elasticity accompanying the slow force decay (Brunello *et al*., 2009). Eventually, in some half-sarcomeres, motors attain the critical level of strain at which rapid detachment occurs (Lombardi and Piazzesi, 1990)promoting the half-sarcomere give, while the half-sarcomeres in which motors are still attached undergo rapid shortening followed by accelerated detachment and completion of the chemomechanical cycle. In this way chaotic relaxation rapidly ends the contraction, a useful requisite for the execution of coordinated movement by antagonist muscles. Given the resistance to stretch of the I-band titin in the ON state, a necessary condition for the chaotic relaxation to occur is that it is preceded by the titin return to the OFF state. This condition is qualitatively verified in Figure 4H by comparing *t*OFF (violet, the parameter that estimates the switching OFF of titin) and *t*_sh_ (green). *t*_OFF_ may be underestimated with respect to that in a muscle contraction, because it is measured in the presence of PNB and thus in the absence of the cooperative effect of myosin motors that slows down the Tm return to the resting position along the thin filament (Caputo *et al*., 1994; Kress *et al*., 1986; Matsuo and Yagi, 2008) and likely the switch OFF of titin.

### Relation with existing *in situ* and *in vitro* studies on titin mechanics

The rise of an elastic element different from myosin motors during muscle contraction was suggested by stiffness measurements early following the stimulation of single fibres from frog skeletal muscle (Bagni et al., 2005; Bagni et al., 2002; Colombini et al., 2010; Fusi *et al*., 2014) and confirmed in both intact (Nocella et al., 2012) and skinned mammalian muscle preparations (Cornachione and Rassier, 2012; Percario et al., 2018). However, a quantitative description of the I-band titin spring *in situ* in the active sarcomere was hampered by the presence, in the A-band in parallel with the I-band titin, of myosin motors which contribute to the mechanical response with their dynamic stiffness (Huxley and Simmons, 1971) and detachment/attachment kinetics (Lombardi and Piazzesi, 1990). Experiments conducted at extremely large SL to suppress the contribution of myosin motors (Leonard and Herzog, 2010; Powers et al., 2014) cannot provide physiologically reliable information. Half-sarcomere stiffness measurements early during isometric force development, when the number of attached motors is small (Powers *et al*., 2020), were interpreted with a complete mechanical model of the half-sarcomere (Pertici et al., 2019) and indicated the presence of an I-band titin spring with an undamped stiffness that at SL >2.7 μm is 6 pN nm^−1^ per htf and independent of SL. The limit of that work was the impossibility to define the titin dynamics, preventing from the quantitative description of the mechanical interface between I-band titin and A-band myosin motors in the time domain relevant for the contraction. To circumvent that problem titin structural dynamics defined with *in vitro* single-molecule mechanics (Kellermayer et al., 1997; Rief et al., 1997; Tskhovrebova et al., 1997) was used. The gap between the undamped I-band titin stiffness and the two orders-of-magnitude smaller static stiffness responsible for the passive force–SL relation was filled introducing the relaxation kinetics of Ig domains’ folding–unfolding (Martonfalvi et al., 2014; Rivas-Pardo et al., 2016). Under these conditions it was calculated that the weak half-sarcomeres lengthened a few hundred nanometres within the few hundred milliseconds of the contraction (Powers *et al*., 2020). Instead, the results in the present paper show that, with the I-band titin in the ON-state, the lengthening necessary to equilibrate an increase in force is ten times smaller (a few tens of nanometres) due to the combination of a 4.2 pN nm^−1^ quasi-instantaneous stiffness and a subsequent slow, viscous-like lengthening with an effective drag coefficient as high as 3.9 pN s nm^−1^. A further limit of the mechanism hypothesised in Powers et al. (2020) becomes evident considering that, at SL within the working range of skeletal muscles (2.1 - 2.7 μm), the extensibility of the I-band titin at rest is substantially based on the straightening out of randomly bent elements of the poly-Ig domain (Linke *et al*., 1998a; Linke *et al*., 1998b; Minajeva et al., 2001), while unfolding of Ig elements becomes relevant in the response to an increase in stress only at SL above 3 μm, at which the contour length of the poly-Ig domain is attained (Granzier et al., 1996; Linke *et al*., 1996; Trombitas *et al*., 1998).

## CONCLUSIONS

A precedently inconceivable efficiency of the half-sarcomere design based on the complementary roles of the A and I band emerges from the present work. The collective myosin motor in the A-band is responsible for the steady production of force and power in isometric and isotonic contractions and for the resistance to forcible lengthening in eccentric contractions; titin in the I-band serves a dual function through a thin filament activation dependent switch between two states: (*i*) at rest (OFF-state), titin exhibits the large extensibility admitted by the randomly oriented elements of the poly-Ig domain to adapt the half-sarcomere length to the physiological range of muscle lengths; (*ii*) following thin filament activation (ON state), at whatever SL the I-band titin acquires the property of a mechanical rectifier: under a pulling force, it efficiently opposes lengthening with a damped elasticity in which the spring shows an effective stiffness of 4 pN nm^−1^ and the viscous-like element an effective drag coefficient of 4 pN s nm^−1^, while under a compressive force it shortens freely as the drag coefficient becomes three orders of magnitude smaller. These properties and the ON-OFF transition kinetics suggest that the molecular switch is the two-thirds proximal segment of the PEVK domain that, following Ca^2+^ activation of the thin filament, binds the nearby actin monomers and that the rectifying mechanism consists in the marked strain sensitivity of the actin-titin interaction. In this way, even at physiological SL, from one side the I-band titin preserves the serially linked half-sarcomeres from the development of length inhomogeneity because any stress undergone by weaker half-sarcomeres for the force excess of the stronger ones is equilibrated with a few % elongation, from the other side it allows unloaded shortening at the maximum velocity accounted for by myosin motors.

This work also opens a new scenario for the understanding of the molecular basis of muscle regulatory mechanisms and related functions. The first detailed quantitative description of the ON-state of the I-band titin and the identification of its control by the Ca^2+^-dependent thin filament activation suggest new *in situ* investigations in demembranated muscle fibres of murine wild type and mutant models on the mechanosensing-based regulation/dysregulation of the signalosomes distributed along the titin domains. Finally, the unexpectedly efficient mechanical link between the Z-end of the half-sarcomere and the A-band titin at physiological SL offers a new basis for understanding the specific role of titin in the mechanosensing-based activation of thick filament (Linari *et al*., 2015).

## ACKNOWLEDGEMENTS

We thank the European Synchrotron Radiation Facility (ESRF) for provision of synchrotron beam time; the staff of the mechanical workshop of the Department of Physics and Astronomy (University of Florence) and Jacques Gorini (ESRF) for electronic and mechanical engineering support; Francesca Corti and Alessia Melani (CeSaL, University of Florence) and the staff of the Animal Facility (ESRF) for animal care. This project was supported by Fondazione Cassa di Risparmio di Firenze (2020.1660); University of Florence (project rictd1819); and the European Synchrotron Radiation Facility. CS was partly supported by the grant European Joint Programme on Rare Diseases 2019 - IDOLS-G.

## AUTHOR CONTRIBUTIONS

Conceptualization, V.L., C.S., G.P., P.B., M.L.; Methodology, V.L., G.P., C.S., P.B.; Formal analysis, G.P., C.S., M.L., M.R.; Investigation, C.S., V.L., P.B.; Resources, T.N., A.M.C; Writing – Original Draft, V.L., G.P.; Writing – Review & Editing, V.L., G.P., C.S., M.R., M.C.,; Visualization, M.C., C.S., P.B., G.P.; Funding Acquisition, V.L., G.P., M.C.

## STAR METHODS

### RESOURCE AVAILABILITY

#### Lead contact

Further information and requests for resources and reagents should be directed to and will be fulfilled by the Lead Contact, Vincenzo Lombardi

#### Materials availability

This study did not generate new unique reagents.

#### Data and code availability

The datasets supporting the current study are available from the corresponding author on request.

### EXPERIMENTAL MODEL AND SUBJECT DETAILS

The experiments were done on single muscle fibres or small fibre bundles (up to 3 fibres each) from *Rana esculenta*, isolated from the tibialis anterior or lumbricalis muscle. Frogs (ca. 50 g weight) were kept at 4-7 °C in the semi-hibernation condition in humid cases that allow them to choose between water and wet environment. Frogs were killed by percussive blow to the head followed by the destruction of the spinal cord in agreement with the Authorization 956/2015-PR from the Italian Health Ministry in compliance with Decreto Legislativo 26/2014 and with EU directive 2010/63. The use of the intact fibre isolated from frog muscle is particularly suitable because (*i*) the parallel elasticity responsible for the passive force-sarcomere length relation is free from the contribution of extra-cellular matrix and solely due to titin (Magid and Law, 1985; Meyer and Lieber, 2018) and (*ii*) mechanical measurements can be conducted by means of a striation follower (Huxley et al., 1981) on a population of sarcomeres selected near the force transducer in order to eliminate inertial effects (Fusi *et al*., 2014). Mechanical experiments have been carried on fibres isolated by tibialis anterior muscle at the PhysioLab, Department of Biology, University of Florence, Florence, Italy. This muscle is preferred to other hind-limb muscles for the quality of dissection and fibre length (*l*_0_, the fibre resting length, that is the length at 2.15 μm SL, is ~5 mm), factors that are relevant for the optimisation of fast sarcomere level mechanics. Combined X-ray diffraction and mechanical experiments have been carried at the ID02 beam line of the European Synchrotron Research Facility (ESRF), Grenoble, France. In these experiments also single fibres or small bundles of lumbricalis muscle were used. Fibres from lumbricalis muscle are shorter (*l*_0_ ~1.7 mm), so that the risk that the lengthening response to a force step saturates the range of movement set in the motor, compromising the X-ray structural response, was minimised. The use of small fibre bundles (two-three fibres) for X-ray diffraction experiments is necessary for a diffracting mass large enough to have a signal-to-noise ratio in the collected X-ray patterns adequate to measure the weaker reflections.

### METHOD DETAILS

#### Preparation and mounting of the fibre

Fibre dissection was performed under a stereomicroscope (Stemi SV6 or SteREO Discovery V8, Zeiss) with the aid of small knives, tweezers and scissors, in a Perspex trough containing Ringer solution, at room temperature. Fibres with the largest diameter were selected during dissection. In this way, given the relation between the histochemical characteristics of twitch fibres of frog muscle and their dimension (Iaizzo, 1990; Spurway and Rowlerson, 1989), it is likely that only type 1 fibres have contributed to this study, as suggested also by the small variation of type-dependent parameter like unloaded shortening velocity (2.53 ± 0.22 μms^−1^ per hs, mean ± SD at 4°C).

Dark-field illumination was used during the dissection. Special care was taken in dissecting and mounting the fibre in order to minimize the amount of tendon compliance in series with the sarcomeres and transverse movements of the fibre during contraction or imposed length/force changes. Fibre tendons were trimmed to a length as short as possible (total length around 300 μm), and held with aluminium foil clips (Ford et al., 1977).

##### Mounting of the fibre for mechanical experiments

The fibre was horizontally mounted in a thermoregulated anodized aluminium trough between the lever arms of a capacitance force transducer (Huxley and Lombardi, 1980) and a loudspeaker motor (range of movement ± 600 μm, upgraded from the original design of (Lombardi and Piazzesi, 1990)) by means of the aluminium clips. The bottom of the trough (in which a channel of the proper dimensions was drilled for the optical path) was covered by a microscope cover-glass. The top of the trough was covered with a cover-glass carrying two platinum plate electrodes running parallel to the fibre. The position of the fibre in the trough and its resting length were adjusted by means of the two micromanipulators carrying the force transducer and the loudspeaker motor. During the experiment the temperature in the trough (4 or 14 °C) was continuously monitored with a thermistor that provided the feedback signal for the temperature control circuit, the output of which fed a thermoelectric module stuck to the bottom of the experimental chamber. The whole system (experimental trough, loudspeaker motor, force transducer) was carried on a metal plate connected to the movable stage of a microscope stand (ACM, Zeiss).

##### Mounting of the fibre for combined X-ray diffraction and mechanical experiments

The fibre bundle was mounted in a thermoregulated trough adapted for X-ray measurements. Two hollow cylinders carrying two mica windows and the stimulating electrodes were moved as close as possible to the fibre, to minimize the X-ray path through the solution. The gap between the windows was typically 600 μm. To have the fibre axis corresponding to the smaller (vertical) size of the X-ray beam and maximize the spatial resolution of X-ray signals along the meridional axis, parallel to the fibre axis, the trough was sealed and mounted vertical at the beamline, with the force transducer on the top and the motor at the bottom.

#### Measurements and stimulation of the fibre

Measurements of sarcomere length (SL), fibre length and cross-sectional area (CSA) were made under ordinary light on the movable stage of the ACM microscope using a 40x water immersion Zeiss objective and a 25x eyepiece. SL was determined by averaging, at different points along the fibre (~ 500 μm apart), the number of sarcomeres in twenty divisions of the graduate scale of the 25x eyepiece. Fibre length was determined by means of a dial gauge mounted on the stand of the ACM microscope, by measuring the displacement of the movable stage in the direction of the fibre axis required to move the two tendon ends of the fibre to the centre of the microscope optic field. Fibre length was initially adjusted to have an average sarcomere length (SL) of 2.15 μm. Different SL (range 2.3-3 μm) were obtained by increasing the fibre length until the desired SL was attained as controlled under the microscope. At the same points at which SL was measured, the height (*h*) and the width (*w*) of the fibre were determined and the fibre CSA was calculated as if the section were elliptical. The height was determined with the fine focusing motion at the middle of the fibre width. The width was read on the graduated scale of the eyepiece.

To elicit the tetanic contraction, a train of even number of stimuli of alternate polarity was applied transversely to the muscle fibre by means of the platinum electrodes across which up to 10 V could be applied with a constant-voltage pulse generator. Stimuli of 1.5 times the threshold and 0.5 ms duration were used. The optimal stimulation frequency, the minimum frequency for a fused tetanus in each fibre at the selected temperature, ranged 18-25 Hz at 4 °C and 50 - 60 Hz at 14 °C and was kept for the subsequent phase of the experiment in the presence of the myosin inhibitor para-nitro-blebbistatin.

#### Solutions

The physiological solution (Ringer) had the following composition: 115 mM NaCl, 2.5 mM KCl, 1.8 mM CaCl2, 3 mM phosphate buffer at pH= 7.1. Para-nitro-blebbistatin, PNB, was dissolved in dimethyl sulfoxide (DMSO, stock solution, 13 mM concentration) and added to the Ringer’s solution to have a final concentration of 20 μM. The addition of DMSO alone to control Ringer at the final concentration (22 mM) as that in the PNB Ringer did not alter the mechanical response of the fibre.

#### Mechanical experiments

##### Mechanical apparatus

The force was recorded by means of a capacitance gauge transducer similar to that described by Huxley & Lombardi (1980). The resonant frequency of the force transducers used in these experiments ranged from 30 to 50 kHz, the sensitivity from 80 to 150 mV/mN and the noise from 2 to 8 mV peak-to-peak.

A striation follower (similar to that described by (Huxley *et al*., 1981)) was used to record the length changes of a population of ~500 sarcomeres selected along the fibre. The instrument measures the longitudinal displacement of the two regions bounding the sarcomere population, each about 10 μm broad and containing six consecutive sarcomeres, by determining the number of sarcomeres that cross the optical fields with interpolation to a precision of about 1% of the striation spacing. The displacements occurring at the level of the two regions are subtracted from each other so that the output signal gives the actual length change undergone by the selected fibre segment. The signal is converted into nm per half-sarcomere on the basis of the average sarcomere length in the segment. The sensitivity of the system is adjusted to 100 mV/nm per half-sarcomere. Systematic errors may result from inhomogeneity of the sarcomere length within the segment and from changes of the sarcomere length in the two regions upon activation. The sarcomere inhomogeneity of the fibres used here was less than 5% at rest and fibres developing gross inhomogeneities on activation were discarded.

The loudspeaker motor was servo-controlled using as a feedback signal the output from either the position sensor on the loudspeaker lever (motor position clamp, *P*m-clamp) or the force transducer (force-clamp). For imposing force steps on the fibre, the motor was first operated in *P*m-clamp and then, at a pre-set time before the force step, was switched to force-clamp mode by a command signal. The return to *P*m-clamp was operated either at a pre-set time by a second command signal or whenever the signal from the loudspeaker position sensor exceeded the values set in a couple of comparators. This procedure provided that the length change required to maintain the force imposed on the fibre could not exceed the range of movement (±600 μm, negative for fibre shortening) within which the motor-length transducer behaves linearly and safely.

The step perturbation in force clamp mode is the most powerful protocol to investigate the structural dynamics of molecular and intermolecular processes because, following the step, the force is kept constant so that the length response of the sarcomeres is not influenced by any length change of in-series compliance and records the molecular transformation under a constant potential energy landscape (Bianco et al., 2014; Rivas-Pardo *et al*., 2016). The main limit to the effectiveness of the method is that, unlike the step imposed in *P*_m_-clamp, for the step in force-clamp the time of propagation of the perturbation from the motor lever through the fibre to the force transducer and the frequency response of the force transducer itself contribute to generate a delay in the feedback loop that determines an upper limit to the gain in the system and consequently to the frequency domain of the perturbation delivered in force-clamp. The low force/stiffness characterizing the passive or PNB treated fibre for the absence of myosin motors further increases the delay of the loop in these experiments, thus increasing the effect of the inertia of the fibre. However, inertial effects are minimized by exploiting the striation follower to record length changes in a population of ~500 sarcomeres near the force transducer end (Fusi *et al*., 2014).

##### Mechanical protocols

At the start of the experiment isometric tetani were elicited under *P*_m_-clamp in normal Ringer solution (control solution) at 4 min intervals with train of stimuli of duration ~300 ms (at 4 °C) and ~260 ms (at 14 °C) at SL 2.15, 2.3, 2.5, 2.7 and 3.0 μm. The order of temperature and SL values was randomly chosen in different experiments. The unloaded shortening velocity (*V*_0_) of the stimulated fibre was measured on the selected population of sarcomeres by imposing, in *P*_m_-clamp, steady shortenings of size and velocity sufficiently large to drop/keep the force to zero. Stepwise rise in force (positive force steps) of amplitude Δ*T* ranging 0.12-0.4 *T*_0,c_ (with *T*_0,c_ the isometric plateau force measured at 4 °C at 2.15 μm SL in control Ringer) were imposed at rest in force-clamp mode to elicit the isotonic lengthening transient in control conditions at different SL. The temperature was then set to 14 °C and the fibre perfused with Ringer solution containing 20 μM PNB. The fibre was stimulated every 5 min to control the progression of the effect of PNB that took 30-40 minutes to be complete. Consistently, the first PNB effect was the progressive inhibition of force with persistence of some degree of half-sarcomere shortening against the end compliance, followed by the disappearance of any half-sarcomere shortening. Positive force steps of 0.12-0.4 *T*_0,c_ were imposed both at rest and during tetanic stimulation at 4 min intervals, to elicit the isotonic lengthening transient at different sarcomere lengths (2.3, 2.5, 2.7 and 3.0 μm). The measurements were done at 4 and 14°C; at either temperature the stimulation frequency and duration was the same as that selected to elicit the fused tetanus in control and the step was imposed at 60 ms following the first stimulus, unless differently specified. The size of the force step and the sarcomere length were set in a random sequence.

In 8 of the 22 fibres used in this work, also the shortening transient in response to a stepwise drop in force (negative force step) was determined either at rest or at different times during stimulation at 3.0 μm SL, at which a restoring force of ~0.1 *T*_0,c_ (or 25 pN per half-thick filament) is present at rest.

#### X-ray Diffraction Protocol

Data in control and in PNB Ringer solution cannot be reliably collected from the same fibre bundle because, given the long time for full force inhibition by PNB, the radiation damage would have progressed so as to affect the later responses. Thus, control and PNB experiments were carried on different preparations. To minimize radiation damage, the beam (~ 300 μm x 50 μm (horizontal x vertical, Full Width at Half Maximum, FWHM), flux of 10^13^ photons/s at a wavelength of ~0.1 nm) was attenuated for bundle alignment and, between X-ray exposures, the trough was vertically translated by 100-200 μm. Two fast electromagnetic shutters in series (tandem shutters) were used to limit the X-ray exposure times to the data collection period.

In control experiments 2D diffraction patterns were first collected with 5 ms time windows at rest and at the plateau of the isometric tetanus at 4 °C, at 2.15 μm. The bundle was slowly stretched to 2.7 μm SL and the protocol repeated. At this SL a 0.25 *T*_0,c_ force step was also imposed at rest and X-ray patterns were collected, with 5 ms time frames, 10 ms before the step and 50 ms after the step (see Figure 2I). The SL of 2.7 μm was chosen because this is the largest SL at which X-ray data can be collected with minimal effects of radiation damage (Reconditi *et al*., 2014), and, on the other side, it is large enough to prevent the lengthening response elicited by a force step of 0.25 *T*_0,c_ at rest to exceed the range of the motor movement (± 600 μm).

In PNB experiments, before starting the perfusion with PNB, the fibre bundle was tetanically stimulated in control Ringer solution at 4 °C and 2.15 μm SL in order to record the force reference *T*_0,c_. The bundle was then perfused with PNB Ringer, the trough was sealed and mounted in the path of the X-ray beam. 2D diffraction patterns were collected at 2.15 and 2.7 μm SL, at 4 °C, with 5 ms time frames at rest and during tetanic stimulation at the same frequency as that used to record the control tetanus at 2.15 μm SL. A force step of 0.25 *T*_0,c_ was imposed on the fibre bundle at 2.7 μm SL both at rest and 60 ms after the first stimulus of the tetanic stimulation and 2D patterns in 5 ms time windows were collected 10 ms before the step and 50 ms (rest) or 2 ms (active) after the step (see Figure 2I). The different timing of the X-ray windows at rest and during stimulation was chosen to isolate the structural change at the end of the rapid elongation phase (phase 2, see below) taking into account the much faster speed of this phase in the stimulated fibre.

#### Mechanical data collection and analysis

Force, motor position and half-sarcomere length changes were recorded with a multifunction I/O board (PXIE-6358, National Instruments). A program written in LabVIEW (National Instrument) was used for signal generation and data acquisition. Data analysis was performed using Excel (Microsoft), OriginPro 8.0 (OriginLab Corporation) and programs written in LabVIEW.

The lengthening (*L*_2_) attained at the end of rapid phase 2 of the transient in response to a positive force step (see Figure S2) is estimated by extrapolating back to the half-time of the step (vertical black lines) the tangent (red line) to the later part (phase 3) of the transient. *V*_3_, the steady velocity of the phase 3 lengthening is estimated by the slope of the tangent. In the stimulated fibre the time after the end of stimulation at which the velocity of lengthening increases as a result of the recovery of the resting condition (*t*_OFF_) is measured at the intersection between the straight lines extrapolated from the slope of phase 3 trace (red line) and that extrapolated from the slope of the subsequent faster lengthening (blue line). Force (pN) per half thick filament (htf) has been calculated from force per CSA, considering a density of thick filaments per CSA equal to 5.87*10^14^ m^−2^ (Appendix A in (Piazzesi et al., 2018)). *L*_2_ and *V*_3_ of the shortening transient following the smallest negative force step (between −8 and −12 pN per htf) imposed at 3 μm SL, were estimated with the same procedure as for the lengthening transient. In the responses to larger negative force steps an early phase 2 was in distinguishable from the later rapid shortening at constant velocity *V*_3_. The *L*_2_-Δ*T* relation and *V*_3_-Δ*T* relation for positive steps were fitted with linear regressions to data pooled from the 22 fibres using a built-in function of Origin Pro software. Given the large and dispersed values of *V*_3_ in response to a quite narrow range of negative force steps (see Figure 3G), in this case the *V*_3_-Δ*T* relation of each fibre was fitted individually and then the slopes and intercepts were averaged.

The time after the first stimulus at which I-band titin switches ON (*t*_ON_) is determined by imposing negative force steps at different times after the first stimulus ( Δ*t*) and measuring the time at which the shortening response deviates from that of the resting fibre. The deviation time is estimated by superimposing, starting from the time of the imposed force step, the shortening responses obtained with different Δ*t* (Figure 4C and D). Plotted in this way, the deviation time decreases by increasing Δ*t* and can be precisely estimated on the difference trace (*DL*) between the resting response and the responses obtained with different Δ*t* (See Figure S3A and B). For each response *t*_ON_ is recovered by adding Δ*t* and the time from the step to the upper deviation of the *DL* trace from the baseline. For instance, in the experiment of Figure 4C, conducted at 4 °C, the response with a stimulus-step interval of 10 ms (orange) deviates from the response at rest at ~4 ms from the step, as marked by the upward deviation from the baseline (black line, representing zero difference) of the orange trace in the *DL* plot (Figure S3A) marks the upward deviation from the baseline. *t*_ON_, calculated by adding 4 ms to Δ*t* (10 ms for the orange response), is 14 ms. With Δ*t* 18 ms (green), the deviation occurs within 1 ms from the step, which is not considered to be a significant delay given the resolution of the system. In this case, though the shortening velocity has not yet attained the steady state active value, the first manifestation of switching ON is assumed to occurs without delay, suggesting that Δ*t* is long enough to be taken as *t*ON and 18 ms is the upper limit for Δ*t*. On the opposite side, reducing Δ*t* to 4 ms the upward deviation of the response (brown) starts at 11 ms from the step, just near the end of the time window afforded in these shortening records, and therefore this Δ*t* is taken as the lower interval limit. The estimate of *t*_ON_ in this case is 15 ms. *t*_ON_ for each of the four fibres used for the experiments at 4°C is estimated by averaging the values measured with 3-7 different Δ*t* (Figure S3C). In the experiments conducted at 14 °C (Figure 4D and Figure S3B), for the deviation to occur within the 10 ms of the shortening time window, the stimulus-step interval must be ~10 ms shorter than at 4 °C. Actually, to obtain a late deviation within the ~10 ms time window (brown), the step must be anticipated with respect to the start of stimulation (negative Δ*t*). The average values of *t*_ON_ at 14 ° for each of the six fibres used (four of which have contributed to data at both temperatures) are reported in Figure S3C.

#### X-ray data collection and analysis

Diffraction patterns were collected on a CCD (Charge Coupled Device) FReLoN (Fast Read out Low-noise) detector with active area 50×50 mm^2^, 2048×2048 pixels and point spread function (PSF) about 80 μm (FWHM). In the experiments described here 8 adjacent pixels along the equatorial direction (perpendicular to the fibre axis) were read as one (8x binning) to reduce the overall readout noise and increase the signal to noise (S/N) ratio without affecting the spatial resolution along the meridional direction (parallel to the fibre axis). Tandem shutter opening and CCD data acquisition were synchronized with the timing of the mechanical protocols by using the same LabVIEW program as that used for mechanical experiments. The intensity of the beam hitting the sample and the time windows of exposures were recorded by collecting a scattered fraction of the beam on a pindiode.

The 2D X-ray patterns were automatically corrected on-line for dark subtraction, flat field and spatial distortion of the detector. The off-line analysis of X-ray data was performed in the laboratory in Florence using Fit2D software (Hammersely, ESRF), PeakFit (SeaSolve Software Inc.) and IgorPro (WaveMetrix Inc.). Single 2D patterns were shifted and rotated to have the four quadrants symmetric relative to the centre of the image using the equatorial 1,0 reflection position, then quadrant folded (mirrored) to enhance the S/N ratio.

A typical 2D pattern from a single fibre from lumbricalis muscle at rest following mirroring is shown in Figure S4. The vertical axis, parallel to the fibre axis, is called the meridian and the reflections along it are the meridional reflections. Those indexed M1 to M6 are orders of a fundamental axial periodicity of ca. 43 nm, associated with myosin. Among them, the M3 reflection from the axial repeat of myosin motors has a periodicity of 14.34 nm (Haselgrove, 1975), and M6, dominated by the periodic mass distribution in the thick filament backbone has a periodicity of 7.17 nm; M1 (which is also contributed from MyBP-C), M2, M4 and M5, the so-called forbidden reflections, signal systematic perturbations of the arrangement of myosin motors on the thick filament. The reflections along the equatorial axis are due to the crystallographic planes originating from the regular disposition of the myofilaments in the lattice. Among them, the most intense are the so-called 1,0 (mainly from the thick filaments) and the 1,1 (from both thick and thin filaments). The off-meridional layer lines parallel to the equatorial axis originate from structures with helicoidal symmetry. Among them, the first myosin layer line reflection at ~43 nm (ML1), is due to the three stranded helical symmetry of the myosin motors on the surface of the thick filament.

The 2D patterns were integrated along the meridian or equatorial axis to get 1D intensity profiles (see Figures 2A–2D) for further analysis. The distribution of diffracted intensity along the meridional axis was determined by integrating 0.012 nm^−1^ on either side of the meridian for comprising and measuring the whole intensity of the reflection and 0.0046 nm^−1^ for measuring their spacings and fine structure with an improved signal-to-noise ratio. The ML1 reflection was integrated in the region between 0.064 and 0.037 nm^−1^ from the meridional axis. The distribution of diffracted intensity along the equatorial axis was calculated by integrating 0.0036 nm^−1^ on either side of the equator.

In the 1D patterns the reflections emerge from a background due to the diffuse scattering from the solution and disordered material present in the preparation. The background intensity distribution was determined using a convex hull algorithm and subtracted (Figures 2B–2D). The 1D intensity profiles of the reflections (with the exception of ML1) were fitted with a Gaussian peak (1,0 and 1,1, integration limits 0.029-0.041 nm^−1^ and 0.052-0.070 nm^−1^, respectively) or multiple Gaussian peaks with the same axial width (M3 and M6, integration limits 0.067-0.072 nm^−1^ and 0.133-0.144 nm^−1^, respectively). The total intensity of a reflection was calculated as the sum of the component peaks and its spacing was determined from the weighted mean of the centres of the component peaks, calibrated using as a reference the position of the M3 reflection in the fibre at rest at full overlap (sarcomere length 2.1-2.2 μm), taken as 14.34 nm.

The intensity of ML1 reflection was obtained by integrating 1D distribution between 0.019–0.023 nm^−1^ corresponding to the half, low angle side of the reflection to avoid contamination with the overlapping, non-resolved AL1 layer line from the actin helix (Piazzesi et al., 1999).

The observed intensity of the reflection is increases with the number of myofilaments in the X-ray beam, which depends on the CSA of the fibre bundle and on possible inhomogeneities along it. It also varies with inverse proportionality to the sarcomere length. To correct for these factors and make the results from different fibre bundles comparable, the intensities of the reflections were scaled by the intensity of the equatorial 1,0 reflection (*I*1,0) at rest as measured on the same pattern at the same SL. The validity of this normalisation procedure depends on the finding that *I*1,0 at rest in different fibres depends solely on the number of myofilaments in the X-ray beam as demonstrated by *I*1,0 remaining intrinsically constant once corrected for the fibre mass under the beam (Reconditi *et al*., 2014).

### QUANTIFICATION AND STATISTICAL ANALYSIS

Data are expressed as mean ± SD or SEM as specified. Mechanical experiments were done on 22 fibres dissected from as many frogs. Given the complexity of the experimental design (combination of force steps of different sizes at different SL and different temperature) not all the fibres contributed all the protocols. The number of fibres contributing to each protocol is reported on the text and in the figure legends. The values of *n* reported for a given parameter refer to the number of repeats contributing to estimate the mean of that parameter.

The total number of preparations used for X-ray experiments was eight from as many frogs. Four bundles were used for measurements in control solution and four for determining the effects of PNB. The values of *n* reported in the legend of Figure 2 refer to the number of repeats of each protocol.

Statistical significance was determined using two-tailed t-test, assuming the level of significance P< 0.02.

