## Supplementary material for "Titin switches from an extensible spring to a mechanical rectifier upon muscle activation": SUPPLEMENTAL INFORMATION 29vii.pdf

### NOTE S1

In this work we assume that the passive force-SL relation of the single fibre from frog skeletal muscle can be explained by the contribution of two spring elements with different extensibility, serially linked in the I-band titin (Linke *et al.*, 1998b; Trombitas *et al.*, 1998): (*i*) the proximal poly-Ig domain that in the SL range 2-2.7 μm behaves as an entropic spring with a large persistence length ($L_p$) and thus responds to a stretch with development of very low force, and (*ii*) the PEVK domain which, at SL ≥ 2.8 μm, at which the poly-Ig spring approaches its contour length ($L_c$) becoming inextensible, responds to further stretch with rise in force according to a much shorter $L_p$. In the absence of direct information on the molecular structure of the frog titin, this explanation of the frog passive force-SL relation relies on the definition of the contributions of the two spring elements in the rat psoas myofibril (Linke *et al.*, 1998a; Linke *et al.*, 1998b). Justifications for the assumption are that (*i*) in either preparation the passive force-SL relation is free from the contribution of the ECM (Meyer and Lieber, 2018), and (*ii*) the titin structure is mostly conserved in ortholog isoforms of the fast skeletal muscle of vertebrates. The assumption is tested here by conducting a comparative analysis of the force – SL relation of the two experimental models (Figure S1 blue dashed line rat psoas myofibril from (Linke *et al.*, 1998a), red dashed line frog muscle fibre from Figure 1B). These two preparations share the same myosin filament length ($l_M$ = 1.6 μm) and therefore the force – SL relation can be uniquely expressed also as force versus the I-band titin length ($l_T$= (SL – $l_M$)/2) (lower abscissa). It can be seen that the two relations exhibit a quite similar large extensibility up to SL ~2.8 μm and then diverge, with the frog fibre relation rising more steeply. This qualitative comparison suggests that (*i*) the poly-Ig entropic elasticity is similar; (*ii*) at SL > 3 μm, when the PEVK elasticity becomes dominant, the frog relation exhibits a larger steepness. For a quantitative test we used the I-band titin model that implies the sequential extension of two serially linked WLC's (Labeit and Kolmerer, 1995; Linke *et al.*, 1998a; Linke *et al.*, 1996; Linke *et al.*, 1998b; Trombitas *et al.*, 1998): the first WLC, provided by the poly-Ig segment, with a purely entropic elasticity and the second WLC, provided by the PEVK segment with entropic and enthalpic elasticities. We found that (*i*) fitting the psoas myofibril relation (blue circles in Figure S1) gives almost the same estimates of the relevant parameters of the model as in the original works (Table in Figure S1), indicating the reliability of our simulation procedure; (*ii*) fitting the frog fibre relation (red circles in Figure S1, reported also in Figure 1B), gave $L_p$ values of the poly-Ig segment and of the PEVK segment similar to those of psoas myofibril, while the Young modulus *E*, characterizing the enthalpic contribution, was 80% larger (Table in Figure S1) accounting for the larger steepness of the force-SL relation. The comparative analysis indicates that in either preparation the entropic elasticity of the poly-Ig domain, attains its limit at ~3 μm SL.

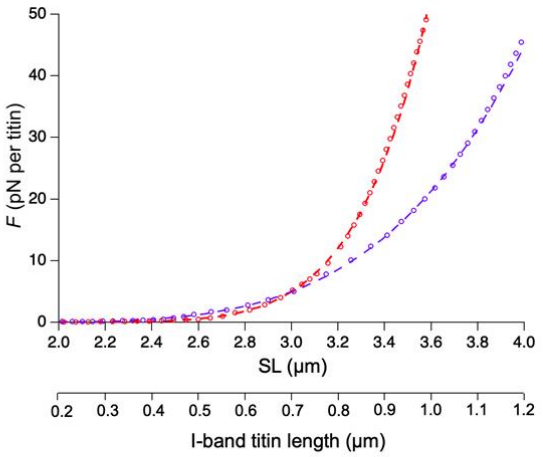


| SL (μm) | Spring element | $L_p$ (nm) | I-band length at $L_c$ (nm) | $E$ (pN) |
|---|---|---|---|---|
| Rat psoas myofibril | | | | |
| < 2.8 | Poly-Ig | 31 ± 2 | 660 ± 10 | - |
| > 2.8 | PEVK | 1.31 ± 0.04 | 1050 ± 80 | 193 ± 6 |
| Frog muscle fibre | | | | |
| < 2.8 | Poly-Ig | 42 ± 1 | 680 ± 30 | - |
| > 2.8 | PEVK | 2.8 ± 0.1 | 910 ± 40 | 331 ± 10 |

**Figure S1. Related to Figure 1. Passive force-SL relation and its simulation by serially linked poly-Ig and PEVK domains.** Dashed lines are the fit to the experimental relations with the empirical exponential equation for the frog muscle fibre (red, from Figure 1B) and for the rat psoas myofibril (blue, from Linke et al., 1998a) with the force expressed in pN per titin molecule. Circles are the result of the simulation obtained assuming the sequential extension of two WLC's, one describing the poly-Ig domain elasticity and the other, integrated by an enthalpic contribution, describing the PEVK domain elasticity. The Table on the right reports the simulation parameters.

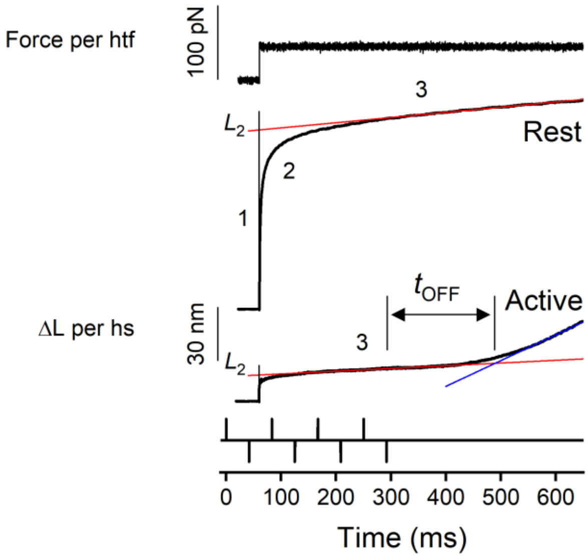


**Figure S2. Related to STAR Methods. Estimate of the relevant parameters of the lengthening transient elicited by a force step.** A force step of 0.2 $T_{0,c}$ (first trace from the top) is imposed on the fibre at rest (second trace) and on the stimulated fibre in the presence of PNB (third trace) at 60 ms following the start of stimulation (fourth trace). The lengthening is composed of an elastic response, phase 1, simultaneous with the force step, followed by a rapid quasi-exponential lengthening, phase 2. Given the speed of phase two and the relatively slow step in force feedback (200-300 μs duration), lengthening in phase 1 merges with the initial part of that in phase 2. Phase 2 lengthening is much smaller and briefer in the active fibre, and in either case is followed by a very slow component (phase 3) at constant velocity. The lengthening attained at the end of phase 2 ($L_2$) is estimated in both the rest and active fibre by extrapolating back to the half-time of the step (vertical black line) the tangent to the linear part of the phase 3 transient (red line). In the active fibre, following the end of stimulation, the velocity of the phase 3 increases again, as a result of the return of titin elasticity to that at rest. The time following the end of stimulation at which the switch occurs ($t_{OFF}$) is measured at the intersection between the straight lines extrapolated from the slopes of phase 3 lengthening (red line) and of the subsequent faster lengthening (blue line).

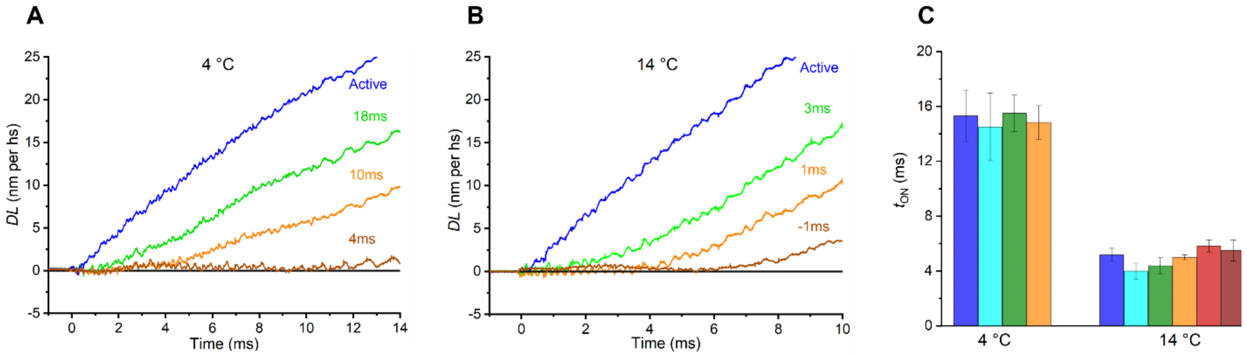


**Figure S3. Related to STAR Methods and to Figure 4C and D. Estimate of $t_{ON}$, the onset of the ON transition.** **A** and **B.** Difference traces (*DL*) obtained by subtracting the shortening response to a negative force step imposed at rest from the responses to the same negative force step imposed at different times ($\Delta t$) after the first stimulus. The traces are superimposed staring from the step time. For each response $t_{ON}$ is recovered by adding $\Delta t$ and the time from the step to the upper deviation of the *DL* trace from the baseline. A refers to the responses at 4 °C in Figure 4C (same color code) and B to those at 14 °C in figure 4D (same color code). **C**. $t_{ON}$ in different fibres (identified by different colours) at the two temperatures, obtained by averaging the values measured with different $\Delta t$. Four fibres (identified by the colour) are contributing to both temperatures.

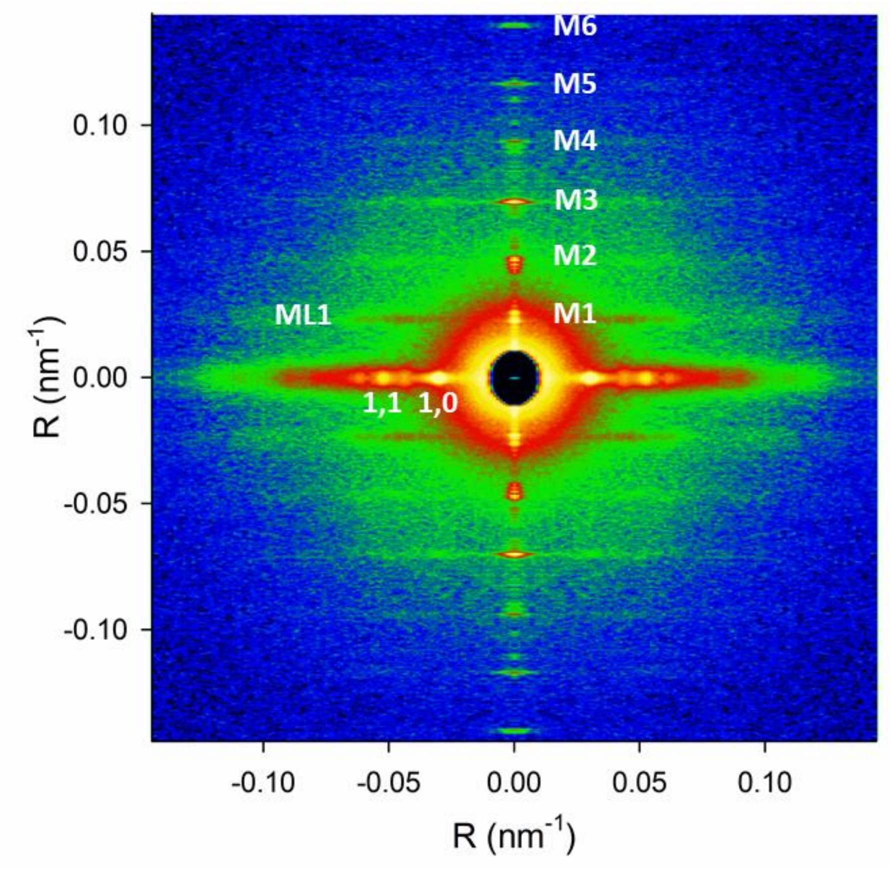


**Figure S4. Related to STAR Methods. 2D X-ray pattern collected from a single muscle fibre at rest in PNB Ringer with a 1.6 m camera length**. SL 2.7 µm, temperature 4 °C. Total exposure time 10 ms. The vertical axis is parallel to the fibre axis.

| SL (μm) | $\Delta T$ (pN per htf) | $L_2$ (nm per hs) | $e_2$ (pN nm$^{-1}$ per htf) | $V_3$ (nm s$^{-1}$ per hs) | $t_{OFF}$ (ms) | *n* |
|---|---|---|---|---|---|---|
| 4 °C | | | | | | |
| Rest | | | | | | |
| 2.7 | 48 ± 3 | 91.61 ± 4.53 | 0.54 ± 0.04 | 22.18 ± 2.15 | | 9 |
| 3.0 | 47 ± 3 | 45.6 ± 3.63 | 1.10 ± 0.09 | 11.30 ± 1.22 | | 14 |
| Active | | | | | | |
| 2.5 | 49 ± 2 | 15.43 ± 0.94 | 3.28 ± 0.16 | 12.35 ± 0.73 | 193 ± 16 | 16 |
| 2.7 | 51 ± 3 | 16.60 ± 0.52 | 3.08 ± 0.11 | 11.38 ± 1.06 | 202 ± 12 | 24 |
| 3.0 | 52 ± 2 | 15.32 ± 1.06 | 3.56 ± 0.22 | 12.73 ± 0.85 | 175 ± 13 | 18 |
| 14 °C | | | | | | |
| Rest | | | | | | |
| 2.7 | 51 ± 4 | 90.82 ± 6.98 | 0.60 ± 0.07 | 27.05 ± 3.07 | | 9 |
| 3.0 | 46 ± 4 | 46.71 ± 4.98 | 1.02 ± 0.08 | 13.03 ± 1.68 | | 7 |
| Active | | | | | | |
| 2.5 | 47 ± 3 | 16.52 ± 1.13 | 2.92 ± 0.28 | 21.05 ± 1.09 | 78 ± 9 | 6 |
| 2.7 | 49 ± 3 | 14.61 ± 0.57 | 3.46 ± 0.26 | 19.53 ± 1.14 | 79 ± 7 | 13 |
| 3.0 | 51 ± 4 | 13.91 ± 1.23 | 3.84 ± 0.33 | 17.15 ± 1.06 | 65 ± 7 | 9 |

**Table S1. Related to Figures 3 and 4. Dependence of the parameters of the lengthening transient on SL at 4 and 14 °C.** The values of the relevant parameters of the lengthening transient (the amplitude of the fast component ($L_2$), the corresponding quasi-instantaneous stiffness per htf ($e_2$) and the velocity of the subsequent slow lengthening ($V_3$)) either at rest or during stimulation (active) are reported as a function of the SL for the ~0.2 $T_{0,c}$ step ($\Delta T$, ~50 pN per htf) at 4 °C and 14 °C. In the active fibre also $t_{OFF}$ (time from the last stimulus for I-band titin to switch OFF) is reported. Data are means ± SEM from 12 fibres (8 fibres (rest) and 12 fibres (active) at 4 °C and 7 fibres (rest) and 11 fibres (active) at 14 °C), *n* is the number of measurements contributing to the average.

| $\Delta T$ (pN per htf) | $L_2$ (nm per hs) | $e_2$ (pN nm$^{-1}$ per htf) | $V_3$ (nm s$^{-1}$ per hs) | $t_{OFF}$ (ms) | n |
|---|---|---|---|---|---|
| **4 °C** | | | | | |
| Rest | | | | | |
| 30 ± 2 | 31.43 ± 5.06 | 1.01 ± 0.15 | 10.49 ± 2.83 | | 4 |
| 47 ± 3 | 45.60 ± 3.63 | 1.10 ± 0.09 | 11.30 ± 1.22 | | 14 |
| 94 ± 11 | 72.00 ± 9.49 | 1.35 ± 0.16 | 17.57 ± 4.38 | | 4 |
| Active | | | | | |
| 31 ± 1 | 8.69 ± 1.12 | 3.82 ± 0.33 | 6.59 ± 0.73 | 171 ± 8 | 15 |
| 54 ± 2 | 15.01 ± 0.67 | 3.76 ± 0.16 | 13.57 ± 0.69 | 167 ± 8 | 33 |
| 111 ± 6 | 32.98 ± 1.87 | 3.42 ± 0.21 | 29.67 ± 4.11 | 162 ± 16 | 10 |
| **14 °C** | | | | | |
| Rest | | | | | |
| 28 ± 3 | 29.94 ± 6.30 | 1.02 ± 0.12 | 9.36 ± 1.49 | | 4 |
| 46 ± 4 | 46.71 ± 4.98 | 1.02 ± 0.08 | 13.03 ± 1.68 | | 7 |
| 95 ± 12 | 58.15 ± 2.42 | 1.65 ± 0.21 | 17.12 ± 4.85 | | 4 |
| Active | | | | | |
| 33 ± 2 | 8.53 ± 1.35 | 4.15 ± 0.35 | 10.94 ± 1.65 | 90 ± 6 | 7 |
| 52 ± 2 | 13.50 ± 0.82 | 3.94 ± 0.22 | 17.55 ± 0.89 | 77 ± 7 | 14 |
| 107 ± 14 | 26.20 ± 2.12 | 4.10 ± 0.53 | 29.03 ± 4.22 | 63 ± 12 | 4 |

**Table S2. Related to Figures 3 and 4. Dependence of the parameters of lengthening transient on the amplitude of a positive force step imposed at 3 μm SL, at 4 and 14 °C.** $L_2$ (amplitude of the fast component of the transient), $e_2$ (quasi-instantaneous stiffness per htf), $V_3$ (velocity of phase 3 lengthening) and $t_{OFF}$ (time after the last stimulus at which ON-OFF transition occurs) are reported for three force step sizes ($\Delta T$): ~0.12 $T_{0,c}$ (~30 pN per htf), ~0.2 $T_{0,c}$ (50 pN per htf) and ~0.4 $T_{0,c}$ (100 pN per htf) imposed both at rest and during stimulation at 4 °C and 14 °C. Data are mean ± SEM from the 22 fibres used in this work, *n* is the number of measurements contributing to the average.

| | Rest | | | Active | | |
|---|---|---|---|---|---|---|
| | Δ*T* (pN per htf) | $V_3$ (nm s$^{-1}$ per hs) | n | Δ*T* (pN per htf) | $V_3$ (nm s$^{-1}$ per hs) | n |
| 4 °C | -12.58 ± 0.60 | -6433 ± 1093 | 12 | -13.19 ± 0.43 | -1780 ± 443 | 17 |
| 14 °C | -12.21 ± 0.52 | -7078 ± 683 | 11 | -12.45 ± 0.91 | -2009 ± 701 | 5 |

**Table S3. Related to Figure 3F. Dependence on activation and temperature of shortening velocity $V_3$ elicited by a negative force step imposed at 3 µm SL.** Data from eight fibres. *n* indicates the number of repeats contributing to each average. Errors are SEM.

| ON transitions | | Low Temperature | | High Temperature | | $Q_{10}$ |
|---|---|---|---|---|---|---|
| $Ca^{2+}$ | $t_{ON}$ | 3.3 [a] | (4°C) | | | 2.5 [b,c] |
| | $t_p$ | 8.5 [d] | (4°C) | | | 2.5 [b,c] |
| AL2 | $t_{ON}$ | 10.5 [e] | (3°C) | | | |
| | $t_{1/2}$ | 16.74 ± 1.05 [e] | (3°C) | 5.85 ± 0.62 [e] | (14°C) | 2.63 [e] |
| | | 12.32 ± 0.34 [e] | (6°C) | | | |
| Titin | $t_{ON}$ | 15.03 ± 0.89 | (4°C) | 4.98 ± 0.22 | (14°C) | 3.02 |
| Myosin | $t_3$ | 15.26 ± 0.38 | (4°C) | 5.87 ± 0.12 | (14°C) | 2.45 |
| OFF transitions | | Low Temperature | | High Temperature | | $Q_{10}$ |
| Titin | $t_{OFF}$ | 191 ± 13 | (4°C) | 70 ± 3 | (14°C) | 2.70 |
| Myosin | $t_{sh}$ | 358 ± 19 | (4°C) | 100 ± 3 | (14°C) | 3.58 |

**Table S4. Related to Figures 4E and 4H. Relevant kinetic parameters accompanying the transitions to the ON state and to the OFF state of the I-band titin and their temperature dependence.** $t_{ON}$, times at which the ON transition of different phenomena starts, $t_{OFF}$, time for the titin OFF transition occurs, $t_p$, time of the peak of $Ca^{2+}$ transient, $t_{1/2}$, half-time of AL2 change, $t_3$, time at which force rises following the first stimulus, $t_{sh}$, time of the transition from isometric to chaotic relaxation. For the data from literature the superscript identifies the reference according to the following list: [a]Sun et al., 1996, [b]Miledi et al., 1982, [c]Eusebi et al., 1983, [d]Caputo et al., 1994, [e]Kress et al., 1986.
